# How does individual trait variation impact the survival of populations with an Allee effect?

**DOI:** 10.64898/2026.03.26.714380

**Authors:** Jonathan Berger, Meike J. Wittmann

## Abstract

The Allee effect is a phenomenon where individual fitness is reduced in small populations, for example because of mate-finding difficulties or increased predation. Allee effects matter in conservation biology because they can drive small populations to extinction. The severity of Allee effects can depend on traits such as mate-search rate and defense against predators. Many natural populations exhibit considerable intraspecific trait variation (ITV) in such traits, but most studies so far assume these traits to be constant. Thus the impact of ITV on populations with Allee effect is largely unknown. Here we create two individual-based stochastic models that simulate a small population experiencing either a mate-finding Allee effect or a predator-driven Allee effect. We analyze how ITV, trait inheritance, and mutation affect the proportion of surviving populations. Under the mate-finding Allee effect, higher ITV hindered population survival and increased Allee thresholds. This can be explained by Jensen’s inequality and the negative curvature of the mate-finding function. Under the predator-driven Allee effect, ITV effects were weak, but higher mutation standard deviations were beneficial, likely because they provided more substrate for selection to act on. We thus recommend to take into account ITV when dealing with threatened populations with an Allee effect.

## Introduction

First described by Allee (1931), the Allee effect describes a scenario where the density or number of conspecifics of an individual has a positive influence on any of its fitness components (Stephens et al., 1999). In cases where such a component Allee effect scales up to the level of total population fitness such that per-capita growth rate is reduced in small populations, we speak of a demographic Allee effect (Fauvergue, 2013; Stephens et al., 1999). In a so-called strong Allee effect, there is a critical population size (Allee threshold) below which populations are expected to decline to extinction (Taylor and Hastings, 2005; Wang and Kot, 2001). Allee effects can be found in many different taxa, including plants, vertebrates, invertebrates, and microbes (Courchamp et al., 1999; Muir et al., 2024). Since Allee effects influence the extinction probability of small populations, they play an important role in conservation biology (Berec et al., 2007) as well as for managing biological invasions and assessing the establishment risk of introduced species (Taylor and Hastings, 2005; Tobin et al., 2011). Allee effects are caused by a variety of mechanisms such as group living, competition, predation, and limitation of mates, resources or pollen (Muir et al., 2024). In this study, we will focus on two types of Allee effect: the mate-finding Allee effect and the predator-driven Allee effect.

A mate-finding Allee effect occurs when an individual’s mating success decreases with decreasing population size (Fauvergue, 2013). Failure to mate can be caused by a variety of mechanisms such as spatial or temporal isolation (Robinet et al., 2007) and rejection of partners in species with mate choice (Bessa-Gomes et al., 2003) and is more likely to occur in solitary species compared to species living in aggregated groups (Dobson and Lees, 1989). Due to the prevalence of sexual reproduction in most plant and animal species, mate-finding Allee effects are thought to be one of the most common types of Allee effect (Robinet et al., 2007). Depending on the circumstances, mate-finding Allee effects can lead to demographic Allee effects (Fauvergue, 2013). Strong evidence of mate-finding Allee effects leading to demographic Allee effects has been found in gypsy moths (*Lymantria dispar*), glanville fritillary butterflies (*Melitaea cinxia*) and the copepod *Hesperodiaptomus shoshone*, with weak or speculative evidence for more species and taxa (Gascoigne et al., 2009; Kramer et al., 2008; Kuussaari et al., 1998; Liebhold and Bascompte, 2003; Sarnelle and Knapp, 2004; Tcheslavskaia et al., 2002).

A predator-driven Allee effect (or more generally a consumer-driven Allee effect, as this phenomenon may occur also in consumer-resource systems e.g., with plants and herbivores) occurs when the predation risk for a prey individual is increased when the prey population size or density is low. In other words, increasing population size or density while small decreases predation risk. This is known as the dilution effect and can arise in different ways. First, for gregarious species, joining together in a bigger group can reduce each individual’s chance to be attacked by a predator (Foster and Treherne, 1981). Banding together in a so called “selfish herd” can even allow individuals in the center of the formation to be completely safe if predator attacks can only come from outside the group (Hamilton, 1971; Morton et al., 1994; Reluga and Viscido, 2005). Second, the dilution effect can also take effect in species that do not show gregarious behavior (Angulo et al., 2007; Wittmer et al., 2005). This happens especially if the predator population can maintain itself off of other prey, has a saturating (type II) functional response, and does not avoid regions of low density of the focal prey species (Angulo et al., 2007; Courchamp et al., 2008; Gascoigne and Lipcius, 2004; Wittmer et al., 2005). Predator-driven Allee effects leading to demographic Allee effects have been proposed in different species including meerkats (*Suricata suricatta*), quokkas (*Setonix brachyurus*), island foxes (*Urocyon littoralis*), caribou (*Rangifer tarandus caribou*), lesser kestrels (*Falco naumanni*) and aphids (*Aphis varians*) (Angulo et al., 2007; Clutton-Brock et al., 1999; Courchamp et al., 2008; Serrano et al., 2005; Sinclair et al., 1998; Turchin and Kareiva, 1989; Wittmer et al., 2005). Thus similar to mate-finding Allee effects, predator-driven Allee effects are likely to play an important role in the extinction of many species (Kramer and Drake, 2010).

Both mate finding and predator-prey interactions are highly trait-dependent. Mate-finding Allee effects are expected to be strongly influenced, for example, by movement parameters such as search rate or search straightness (Berec, 2018; Fagan et al., 2010), mating system (monogamy vs. polygamy, Bessa-Gomes et al., 2003), choosiness of females (Bessa-Gomes et al., 2003), gamete parameters such as sperm swimming speed and longevity for free-spawning marine animals (Levitan, 2000), or the ability or propensity to self-fertilize (Kramer et al., 2009). Predator-prey interactions can be influenced by morphological and behavioral traits of both predator and prey individuals (Hammill et al., 2023; Kreuzinger-Janik et al., 2019; McCoy and Bolker, 2008; Milonas et al., 2011; Toscano and Griffen, 2014). As an example for a morphological trait, both the prey’s and predator’s body size can impact handling times and functional response of the predator (Hammill et al., 2023; Kreuzinger-Janik et al., 2019; McCoy and Bolker, 2008; Milonas et al., 2011). Behavioral traits like activity levels are another factor that can change interactions and survival chances in predator-prey systems (Toscano and Griffen, 2014).

The morphological and behavioral traits affecting mate finding and predator-prey interactions often exhibit considerable intraspecific trait variation (ITV). For example, in some sea urchin species intraspecific variation in egg size and sperm swimming speed affects fertilization success (Levitan, 1996, 2000). In hermaphroditic plants and animals, there can be intraspecific variation in the ability or propensity to self-fertilize (Cheptou and Avendaño V, 2006; Ramm et al., 2012). In many prey or resource species, there is variation in defense levels (Bolnick et al., 2011; Kawagoe et al., 2011; Moore et al., 2014; Speed et al., 2012). These differences between individuals within a species arise from genetic variability, acclimatization, or phenotypic plasticity (Albert et al., 2011; Kawagoe et al., 2011; Kuppler et al., 2020). So far, most ecological models with an Allee effect are based on population mean trait values and ignore ITV (Boukal and Berec, 2002; Courchamp et al., 2008). However, ITV can have important ecological consequences at the population, community or ecosystem level (Bolnick et al., 2011; Moran et al., 2016; Westerband et al., 2021).

The ecological consequences of ITV arise via a number of mechanisms, including nonlinear averaging (Jensen’s inequality), the portfolio effect, and eco-evolutionary dynamics (Bolnick et al., 2011). Nonlinear averaging is the phenomenon that the average of a nonlinear function of a variable trait is in general not the function value of the average trait in the population. Jensen’s inequality (Denny, 2017; Jensen, 1906; Ruel and Ayres, 1999) states more specifically that if the function has a positive curvature (second derivative), the effect of ITV on the average function value will be positive, and if the function has a negative curvature, the effect of ITV will be negative. Via nonlinear averaging, ITV can for example promote or hinder coexistence in competition and predation scenarios (Gibert and Brassil, 2014; Hart et al., 2016; Lichstein et al., 2007; van Benthem et al., 2025). The portfolio effect of intraspecific variation is the phenomenon that different responses of individuals, subpopulations, or strains in a population to fluctuating environmental conditions can buffer overall population dynamics (Bolnick et al., 2011; de Bruin et al., 2024; Schindler et al., 2010). If ITV has a heritable basis, it can also affect eco-evolutionary dynamics (Hendry, 2017), which could for example lessen the consequences of habitat loss (Bagawade et al., 2024). Because of these various and pervasive effects of ITV on many ecological processes, we hypothesize ITV to also affect the dynamics of populations subject to Allee effects.

So far, there are just a few isolated results on the consequences of ITV for Allee effects. Although in one of the first papers on the Allee effect, Dennis (1989) already showed how the average mating probability can be computed if mate-search rate is not constant across the population, but varies according to some probability distribution, he did not investigate the consequences of different levels of ITV. Berec (2018) considered individual variation in mate search rate or search straightness. He found that ITV in mate search rate when males are the searching sex does not have an effect, but ITV in mate search rate when females are the searching sex reduced female mating probability. ITV in search straightness increased female mating probability. This raises the question why different kinds of ITV have different effects on mating probability and what the consequences for population persistence and Allee thresholds will be. The most detailed investigation of the consequences of ITV for Allee effects so far investigated individual variation in timing of reproduction (reproductive asynchrony). These models suggest that the more the individual reproductive periods are spread out over the time, the lower the probability for females to successfully mate at low density and the higher the Allee threshold (Calabrese and Fagan, 2004). However, when there is also protandry or protogyny (where the mean maturation date differs between sexes) sometimes higher or intermediate levels of ITV in maturation date can also be optimal for mating success because they can increase overlap in active periods between the sexes (Larsen et al., 2013; Robinet et al., 2007).

Natural populations are expected to exhibit ITV in many traits that can potentially affect Allee effects. In this study, we thus create individual-based stochastic models that incorporate heritable or non-heritable ITV in traits influencing either a mate-finding Allee effect or a predator-driven Allee effect. Our goal is to examine how intraspecific trait variation and inheritance affect the survival probability of a small founder population with an Allee effect and to elucidate under what conditions ITV is expected to alleviate or worsen Allee effects.

## Model and simulations

### Model description

For both the mate-finding Allee effect model and the predator-driven Allee effect model, a founder population of size *N*_0_ is generated and each individual gets assigned a sex (male or female) with equal probability (see Table 1 for an overview of all parameters). Furthermore, the trait value *z* for each individual is drawn from a Gamma distribution with mean trait value *µ* and trait standard deviation *σ*. That is, we have a probability density function

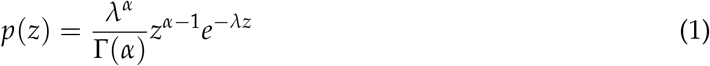

where the shape parameter

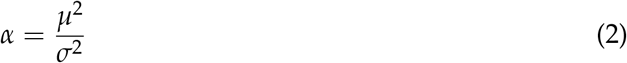

and the rate parameter

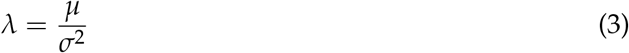

are chosen to produce the desired mean and standard deviation. We used a Gamma distribution rather than a normal distribution so that traits can take only positive values (trait space is (0, ∞)), which is realistic for the types of trait we will be considering.

**Table 1.** List of all the parameters and variables that were used in either the mate-finding or predator-driven Allee effect model.

| Parameter/Variable | Explanation | Default value |
| --- | --- | --- |
|  | <u>Mate finding</u> |  |
| $z$ | Individual mate-search rate | 0.02 |
|  | <u>Predator-driven</u> |  |
| $z$ | Individual defense trait | 0.02 |
| $P$ | Number of predators | 1 |
| $c$ | Maximum attack rate of the predator | 2 |
| $b$ | Decay rate of the attack rate with increasing defense trait value | 70 |
| $h$ | Handling time of the predator | 0.03 |
|  | <u>Both models</u> |  |
| $n_{\max}$ | Number of generations simulated | 100 |
| $N_0$ | Founder population size | 500 |
| $K$ | Carrying capacity of the habitat | 5000 |
| Female proportion | Chance an individual is a female | 0.5 |
| $w$ | Fecundity/average number of offspring per mated female | 2.1 |
| $\mu$ | Mean of the initial trait distribution | 0.02 |
| $\sigma$ | Standard deviation of the initial trait distribution | 0.005 |
| $\sigma_{\text{mut}}$ | Mutation standard deviation | 0.0005 |
| Replicates | Number of replicates | 500 |

After initialization, we simulate the population for *n*_max_ generations. Each generation consists of a life cycle with two steps, but the two model versions differ in the trait-dependent first step. In the mate-finding Allee effect model, the first step is mate finding. In the predator-driven Allee effect model, the first step is predation. The separate first steps are described in the following two paragraphs. The second step in both models is reproduction and is described afterwards.

In the mate-finding Allee effect model, the trait *z* represents an individual female’s mate-search rate. Individuals could differ in their mate-search rate for example because of different movement rates, different abilities to detect nearby males, or different degrees of choosiness (Berec, 2018; Bessa-Gomes et al., 2003). A female with trait *z* meets a male mating partner with probability 1 − *e*^−*z*·*M*^, where *M* is the current number of males. This expression is used in many models of mate-finding Allee effects (McCarthy, 1997; Wittmann et al., 2018) and provides a good approximation to mate-finding probabilities in individual-based models where movement and encounters of males and females in space are explicitly modeled (Berec, 2018). An Allee effect emerges in this model because smaller populations have fewer males, leading to a smaller mate-finding probability.

In the predator-driven Allee effect model, *z* represents a trait that reduces the attack rate of the predator. Such traits could be behavioral defenses like reduced movement or predator avoidance that vary in their effectiveness (Hammill et al., 2015) or morphological or chemical defenses, which also often exhibit a lot of intraspecific variation in both plants and animals (Moore et al., 2014; Sato, 2018; Speed et al., 2012). But the trait could also just be body size if larger prey individuals are harder to attack. Following McCoy and Bolker (2008), we assume that the predator’s attack rate *a* declines exponentially with increasing prey defense trait value:

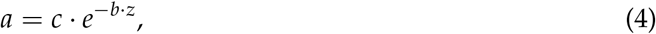

where *c* is the highest possible attack rate, which occurs when the prey is undefended. The parameter *b* regulates the decay of the attack rate with increasing trait value *z*.

The predators in the predator-driven Allee effect model exhibit a type II functional response. That is, if all prey had equal trait values leading to an attack rate *a*, the number of prey individuals consumed per predator per time unit would be

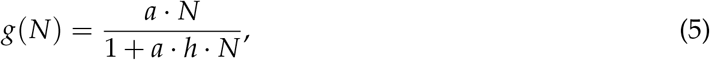

with *h* being the handling time and *N* the total number of prey individuals (Holling, 1959). In order to incorporate prey trait variation and calculate the chance that any specific prey individual gets captured and eaten by a single predator, the following function is used:

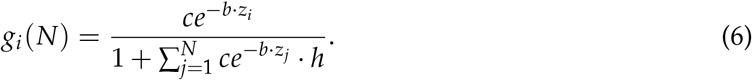

In this case, the survival of a prey individual is not only determined by its own trait *z*_*i*_ that reduces the attack rate of the predator but depends also on the traits of the other prey individuals *z*_*j*_ in the population because these trait values influence how many other prey the predator catches and takes time to handle. In order to get the survival probability *f*_*i*_ of a prey individual *i* as a function of predator population density, its capture probability is multiplied with the density of predators *P* and subtracted from 1:

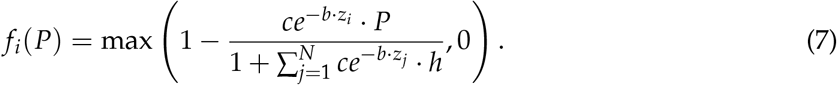

Here we use the maximum to prevent the survival probability from falling below 0. This equation also shows how the Allee effect arises. The survival probability increases with the number of prey individuals because handling other prey individuals takes up the predators’ time and reduces the time for hunting the focal prey individual. Each individual *i* then randomly survives with probability *f*_*i*_(*P*). All the surviving females will mate, with the mates being drawn from the surviving males. As long as there is still at least one male and one female present, there are no mate-finding problems in the predator-driven Allee effect model.

In both models, for each mating female that survived and mated, one random male mate gets drawn to sire all her offspring. The chance to be drawn is equal for all males and males can mate with an unlimited number of females. The number of offspring of each mated pair is drawn from a Poisson distribution with mean *w*, the fecundity parameter. We assume *w >* 2 because for *w <* 2, a population would be expected to go extinct even without an Allee effect.

We assume a simplified inheritance system where each offspring inherits the mean value of their parents’ trait values. Additionally, the inherited trait value is mutated. Using a normal distribution with a mean of zero and standard deviation *σ*_mut_, a number is drawn for each offspring. This number is added to the original trait value of the offspring. In cases where the trait value would fall below 0 it gets replaced with 0. Afterwards each offspring individual gets assigned a sex, again with a 50:50 sex ratio. The parent generation then dies and is completely replaced by the offspring generation. If the new population size exceeds the carrying capacity, randomly chosen individuals are removed from the population until the population size is equal to the carrying capacity.

To investigate the effects of ITV and inheritance, in addition to the standard model (“ITV + Inheritance”), we created modified versions of both the mate-finding Allee effect model and the predator-driven Allee effect model. In the “Inheritance only” model version, intraspecific variation is turned off, which means that there is no variation in the founder generation such that every individual shares the set trait value. The “ITV only” model version turns off both inheritance and new mutations. Instead, offspring individuals get assigned a trait value drawn from a Gamma distribution with mean *µ* and standard deviation *σ*, as for the founder population. Finally, the “None” model version omits both intraspecific variation and inheritance. In practice this means that all individuals from the founder generation share the same trait value and all offspring are assigned that value as well.

### Simulations

In order to assess the effect of ITV and inheritance on the survival of a population for the two models, each of the model versions (ITV + Inheritance, ITV only, Inheritance only, None) are run with various starting population sizes. The rest of the parameters are kept at their standard value (see Table 1). Each simulation is run for *n*_max_ generations (100). After the last timestep, we check whether or not the population survived, meaning whether any individuals are still alive. This is repeated for the set number of replicates. For each founder population size, the proportion of surviving populations over all replicates is calculated. For further analysis, the same process is repeated using different values for standard deviations of the trait and mutations. These parameters are of interest here since they impact how much ITV exists in the population. The simulations and analysis were performed using R (version 4.3.2, R Core Team, 2023).

### Analytical calculation of the Allee threshold

For the “ITV only” model version without inheritance, we can analytically calculate or at least approximate the Allee threshold and how it is affected by ITV. For these analyses, we make the assumptions that at the Allee threshold, the population is still large enough so that the sex ratio does not deviate substantially from 50:50 and that the actual trait distribution in the population is close to the theoretically assumed distribution. That is, we assume that sampling effects of small population size are not playing a large role.

In models with a strong Allee effect, the Allee threshold is an unstable equilibrium of the population model. That is, at the Allee threshold, the expected population size in the next generation is equal to the current population size, whereas for larger population sizes population size is expected to grow and for smaller population sizes population size is expected to decline. Thus, the general idea of computing the Allee threshold is to compute the expected population size in the next generation given that the population currently has size *N*^∗^ and set it equal to *N*^∗^. Then we can solve this equation for *N*^∗^. Here we do not include density regulation because we consider only dynamics at small population density and, given the functions we chose, the expected population size in the next generation will monotonically increase with increasing population size because of higher mating rates or lower predation rates. Thus, there will only be one solution for *N*^∗^, which is the Allee threshold. If *N*^∗^ is larger than the carrying capacity, *K*, the population would always go extinct. Taking the derivative of *N*^∗^ with respect to the trait variance will then inform us about the effects of ITV. We will now perform this analysis separately for the mate-finding Allee effect and for the predator-driven Allee effect.

#### Mate-finding Allee effect

To find the Allee threshold for the mate-finding Allee effect, we have to solve

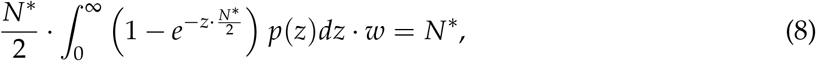

where the left-hand side is the expected population size in the next generation for a current population size of *N*^∗^. The first *N*^∗^/2 represents the number of females and the *N*^∗^/2 in the exponential represents the number of males. The integral represents the average mating success of females with different trait values *z* in the population. Substituting (1), integrating, and solving for *N*^∗^ using the Maxima software (Maxima, 2024, see Notebook and corresponding html file in the Supplementary materials), we obtain

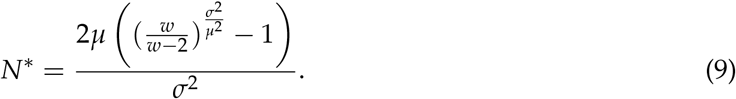

To check whether *N*^∗^ decreases or increases with ITV, we treat *σ*^2^ as the symbol for trait variance and compute

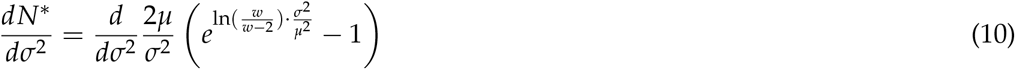

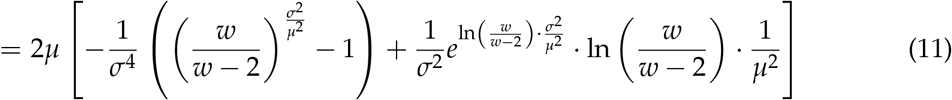

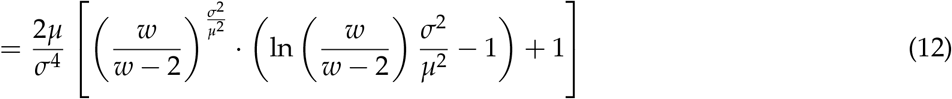

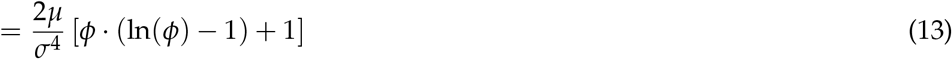

with 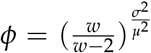. Thus the Allee threshold increases with ITV if *ψ*(*ϕ*):= *ϕ* · (ln(*ϕ*) − 1) + 1 is positive. Since *w > w* − 2, *ϕ >* 1. Since *ψ*(1) = 0 and *ψ*^′^(*ϕ*) = ln(*ϕ*) *>* 0 for *ϕ >* 1, *ψ*(*ϕ*) is positive for all *ϕ >* 1 and thus 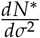 is always positive. Thus any increase in ITV leads to a higher Allee threshold and thus worsens this type of mate-finding Allee effect.

To find the Allee threshold for the extreme case without ITV, one can either use L’Hôpital’s rule to take the limit of (9) as *σ*^2^ goes to 0 or, analogously to (8), solve

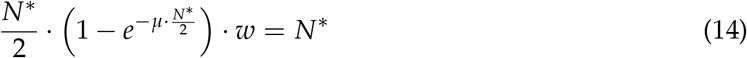

for *N*^∗^. In both cases, we obtain

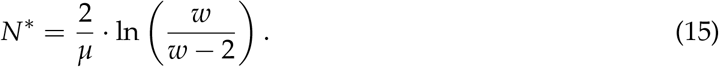

#### Predator-driven Allee effect

To find the Allee threshold for the predator-prey model, we have to solve

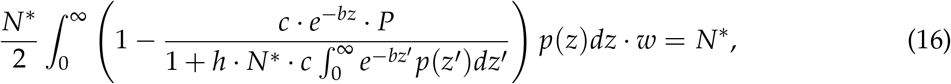

where the left-hand side again represents the expected population size in the next generation if the current population size is *N*^∗^. Let *A* = exp(−*bµ*) without ITV, i.e. with *σ*^2^ = 0, and otherwise

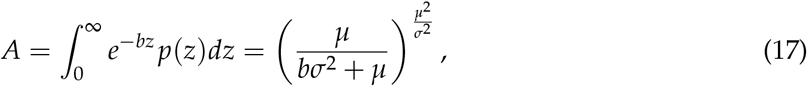

where the integral over the Gamma distribution was again computed using Maxima (2024). Then (16) is equivalent to

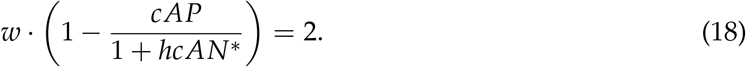

Solving for *N*^∗^, we obtain

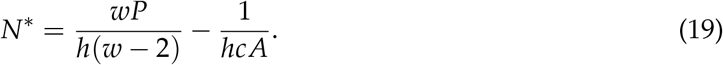

Since

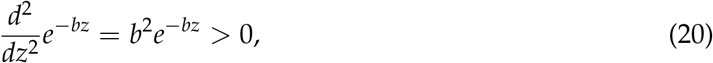

Jensen’s inequality tells us that ITV will increase the average value *A* of *e*^−*bz*^ across the population. Also, to see that *A* is monotonically increasing with *σ*^2^, treat *σ*^2^ again as a symbol for trait variance and consider

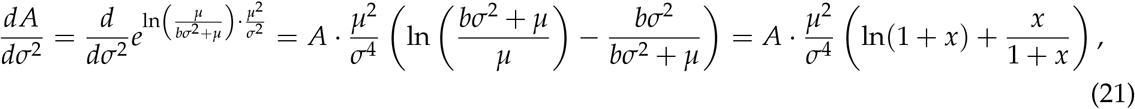

with 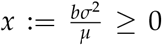. Thus the direction of the effect of *σ*^2^ on *A* depends on the sign of 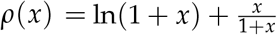. Since *ρ*(0) = 0 and

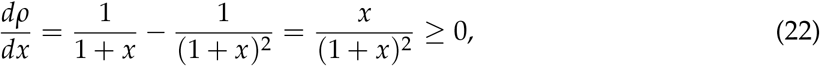

*A* is monotonically increasing with *σ*^2^.

Inspecting (19), we see that an increase in *A*, leads to an increase in *N*^∗^. Thus under the predator-driven Allee effect, ITV also always increases the Allee threshold and worsens Allee effects. However, note that this result is entirely driven by the curvature of the attack rate function *e*^−*bz*^. With this function, attack rate declines quickly when the trait *z* increases. A biologically similar model where the attack rate first decreases more slowly and then falls more rapidly, i.e. a function with a negative curvature, would have given the opposite result that ITV reduces the Allee threshold.

## Results

### Mate-finding Allee effect

With the standard parameters, all four versions of the mate-finding Allee effect model (ITV and inheritance, ITV only, inheritance only, none) show a sigmoid relationship between founder population size and the proportion of surviving populations (Fig. 1 A), as expected for populations with an Allee effect (Dennis, 2002). The success probability curve for the model version that only includes ITV has the same shape as the curves for the other scenarios, but starts to increase at a slightly higher founder population size and reaches its plateau a bit later than the other three model versions (Fig. 1 A). The analytically calculated Allee thresholds (indicated by vertical lines) are approximately where the success probability curves for the individual-based models have their inflection point (the point of steepest increase), as expected based on the theory by Dennis (2002). That is, the simplified analytical calculations give useful guidance regarding the behavior of the full model.

**Figure 1.**
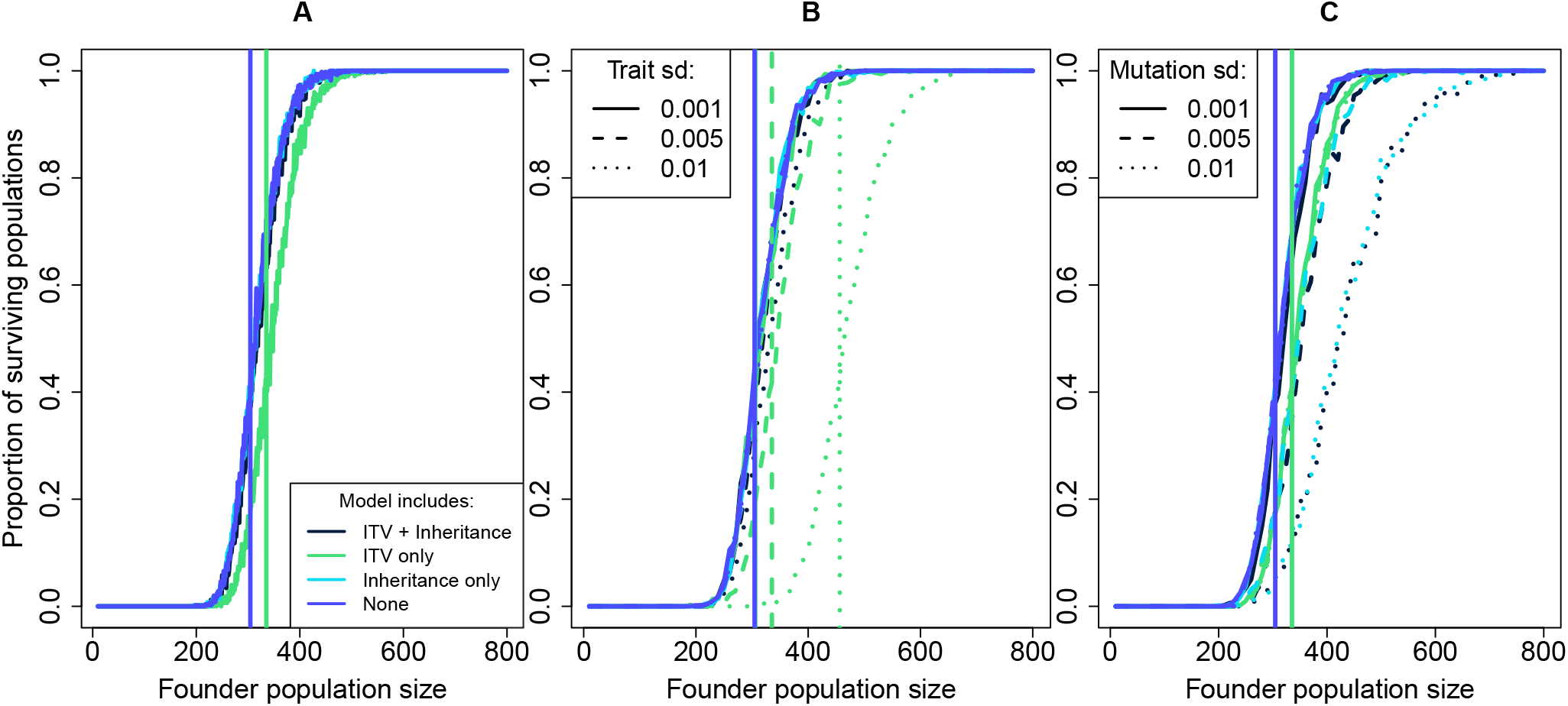
Proportion of surviving populations as a function of founder population size under the four different mate-finding Allee effect model versions with 500 replicates per founder population size. The vertical lines indicate the respective Allee thresholds calculated via the analytical approximation. A) Results for the default parameter settings from Table 1. Here, we ran simulations for all founder population sizes from 10 to 800 (steps of 1). B) Results for different values of the trait standard deviation *σ*. C) Results for different values of the mutation standard deviation *σ*_mut_. In B and C, we simulated founder population sizes between 10 and 800 in steps of 10.

An increase in the trait standard deviation *σ* shifted the success probability curves to the right (that is, caused higher Allee thresholds) for the model versions with ITV and inheritance and especially for the model version with only ITV (Fig. 1 B). For the other two model versions, there was no ITV and thus the trait standard deviation did not have an effect.

Similarly, an increase in the mutation standard deviation shifted the success probability curves to the right and thus caused higher Allee thresholds in those model versions that included mutation (ITV + inheritance and inheritance only, Fig. 1 C). The success probability curves for these two model versions with inheritance and mutations are similar for all values of the mutation standard deviation (Fig. 1 C). For mutation standard deviation 0.005, they are also similar to the success probability curve with only ITV, while for a mutation standard deviation of 0.001 they are similar to the curves of the model that includes neither ITV nor inheritance (Fig. 1 C).

Fig. 2 shows example dynamics of population size (A), population mean trait value (B), and trait standard deviation (C) for the model version with ITV and inheritance, an initial population size of 500, and three different mutation standard deviations (see Fig. 1 B for the corresponding population survival probabilities). Populations with the two smaller mutation standard deviations evolved very little (Fig. 2 B), but grew from the start with a logistic-like sigmoid population size trajectory (Fig. 2 A). By contrast, average population size with the largest mutation standard deviation declined initially before recovering (Fig. 2 A), although many individual replicates either grew or declined from the start or first lingered around the starting population size before increasing (see Fig. S1). These populations with the largest mutation standard deviation showed the strongest evolutionary response (Fig. 2 B). The average trait standard deviation decreased over time for the smallest mutation standard deviation, stayed roughly constant for the intermediate mutation standard deviation and increased for the largest mutation standard deviation, in each case reaching an equilibrium after roughly 20 generations (Fig. 2 C).

**Figure 2.**
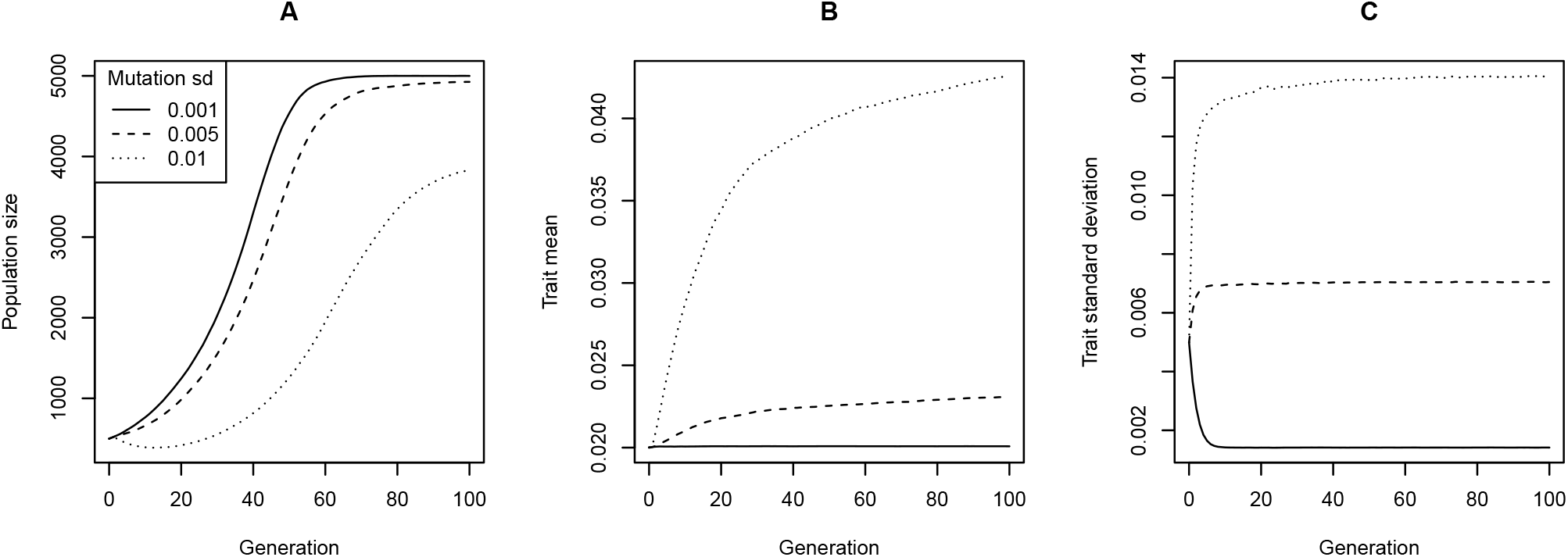
Eco-evolutionary dynamics of the mate-finding Allee effect model with ITV and inheritance and different mutation standard deviations. A) Mean populations size over 500 replicates (counting extinct populations as 0). B) Corresponding mean trait value across all populations that are not extinct at the given generation. C) Average trait standard deviation across all non-extinct replicates. The initial population size was 500 and all other parameters were at their default values from Table 1.

### Predator-driven Allee effect

In the predator-driven Allee effect model with the standard parameter settings, the success probabilities did not differ substantially between the four model versions (Fig. 3 A) or between different values of the trait standard deviation (Fig. 3 B). However, higher mutation standard deviations led to higher success probabilities and smaller Allee thresholds under the model versions with inheritance (ITV + inheritance or only inheritance) (Fig. 3 C). For each value of the mutation standard deviation, both these model versions had similar curves (Fig. 3 C).

**Figure 3.**
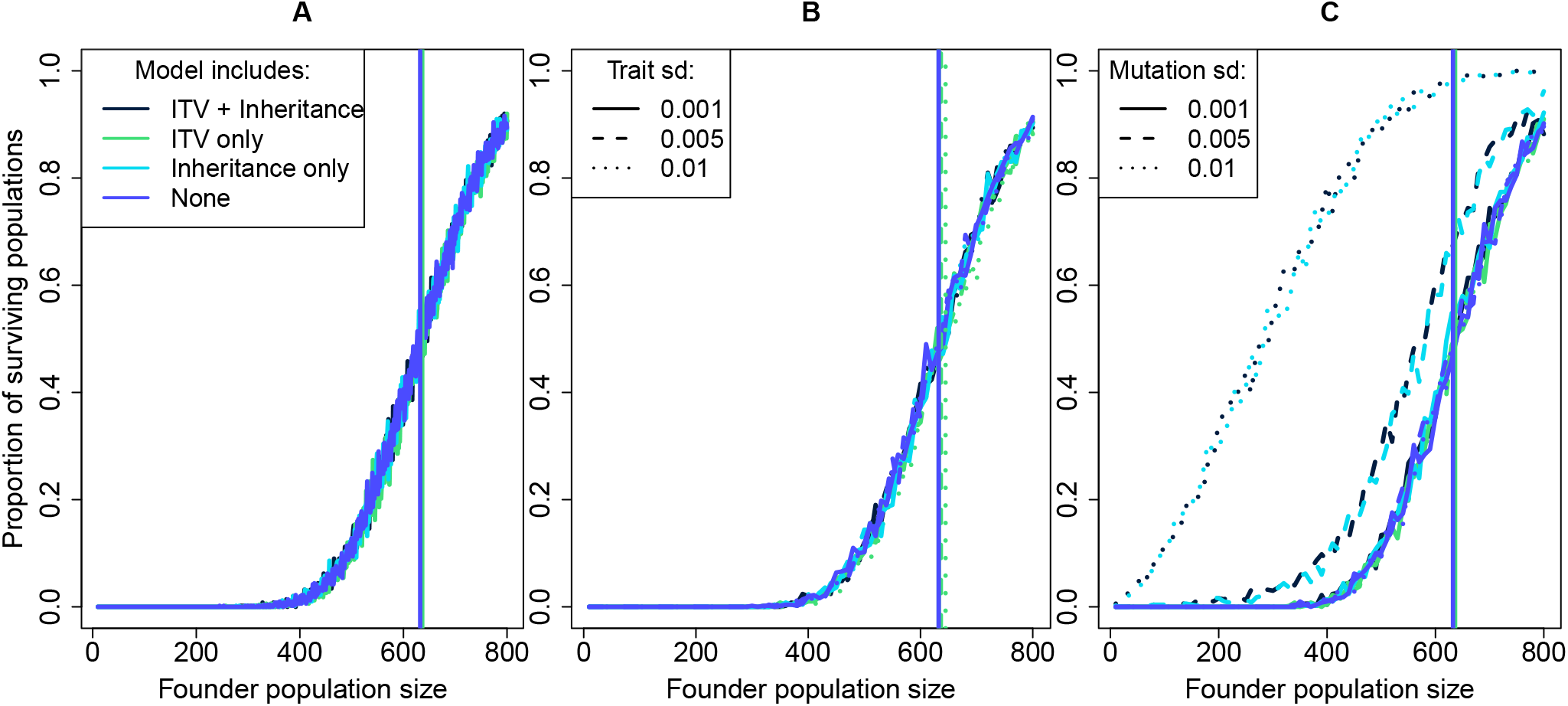
Proportion of surviving populations of the four different predator-driven Allee effect model versions for different founder population sizes with 500 replicates each. The vertical lines indicate the respective Allee thresholds calculated via the analytical approximation. A) Results for the default parameter settings from Table 1. Here, we ran simulations for all founder population sizes from 10 to 800 (steps of 1). B) Results for different values of the trait standard deviation *σ*. C) Results for different values of the mutation standard deviation *σ*_mut_. In B and C, we simulated founder population sizes between 10 and 800 in steps of 10.

Example dynamics for an initial population size of 500 under the different mutation standard deviations are shown in Fig. 4. The mean population size across replicates for all three values dropped at the start, but then started to increase after about 50 years (Fig. 4 A). This increase was the earliest and sharpest and led to the highest mean population size for the mutation standard deviation value of 0.01 (Fig. 4 A, see Fig. S2 for trajectories of individual replicates). While a mutation standard deviation of 0.001 led to almost no increase in the mean trait value (Fig. 4 B), mutation standard deviations of 0.005 and especially 0.01 led to more substantial increases in the mean trait (Fig. 4 B). The dynamics of the trait standard deviation (Fig. 4 C) were similar qualitatively and quantitatively to those for the mate-finding Allee effect model (see Fig. 2 C): the level at which the trait standard deviation equilibrated increased with the mutation standard deviation.

**Figure 4.**
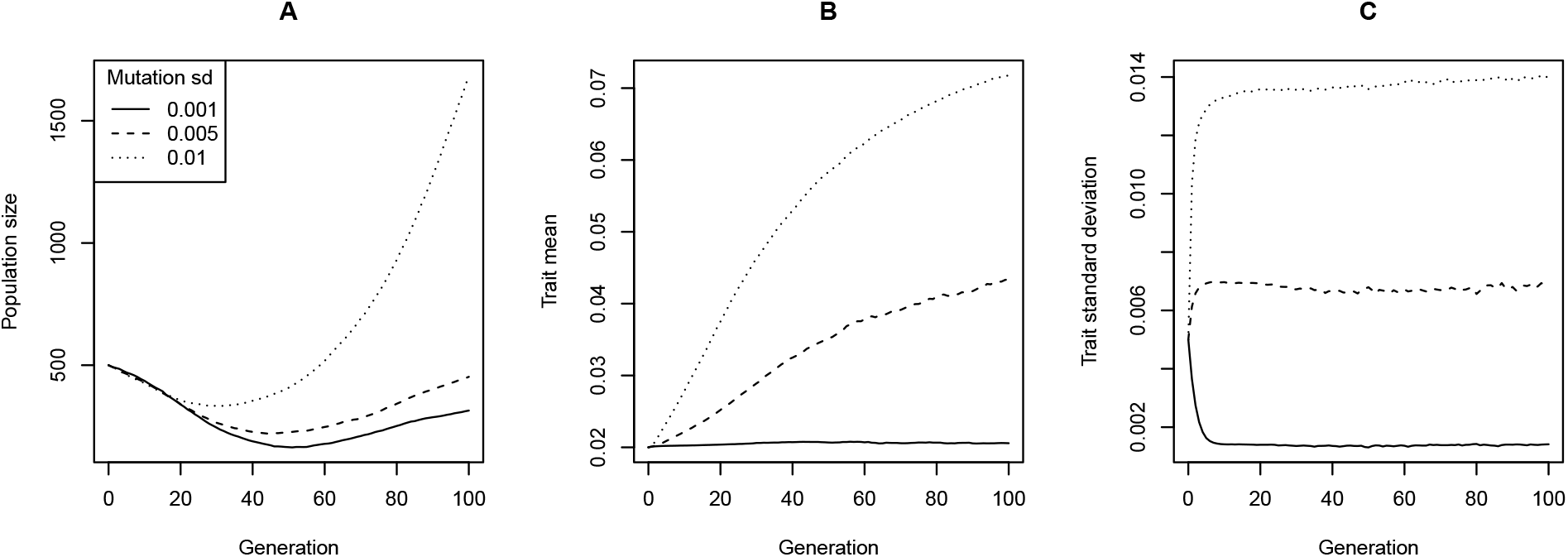
Eco-evolutionary dynamics of the predator-driven Allee effect model with ITV and inheritance and different mutation standard deviations. A) Mean populations size over 500 replicates (counting extinct populations as 0). B) Corresponding mean trait value across all populations that are not extinct at the given generation. C) Average trait standard deviation across all non-extinct replicates. The initial population size was 500 and all other parameters were at their default values from Table 1.

### Estimates of the Jensen gap

For both the function used to calculate the mating success in the mate-finding Allee effect model as well as the function used for the survival probability in the predator-driven Allee effect model, we estimated the “Jensen gap” (Giménez and Jenkins, 2024), that is the change in mating probability or survival probability due to nonlinear averaging of ITV. For both functions, the function value was plotted against the inserted trait value (Fig. 5). The 1000 inserted trait values were drawn from a Gamma distribution using the standard mean trait value and standard trait standard deviation (see Table 1). In case of the survival function, for each of the 1000 data points independently, another 100 trait values were drawn from the same distribution to represent the rest of the population (for the sum in the denominator of Eq. 7). The proportional change due to ITV was calculated as the difference between the average function outcome and the outcome for the average trait value, divided by the outcome for the average trait value (Fig. 5 A). When computing the outcome for the average trait value for the predation survival function, we assumed that the rest of the population all shared the average trait value to truly represent a population without ITV (Fig. 5 B). Both functions have a similar curvature and result in a negative estimate of the effect of ITV (Fig. 5). However, for the predation survival function, the negative estimate of the effect of ITV is weaker compared to the mate finding function (Fig. 5). The “background noise” due to variation in the traits of the 100 other individuals explains why the points in B are not exactly on one curve as in A.

**Figure 5.**
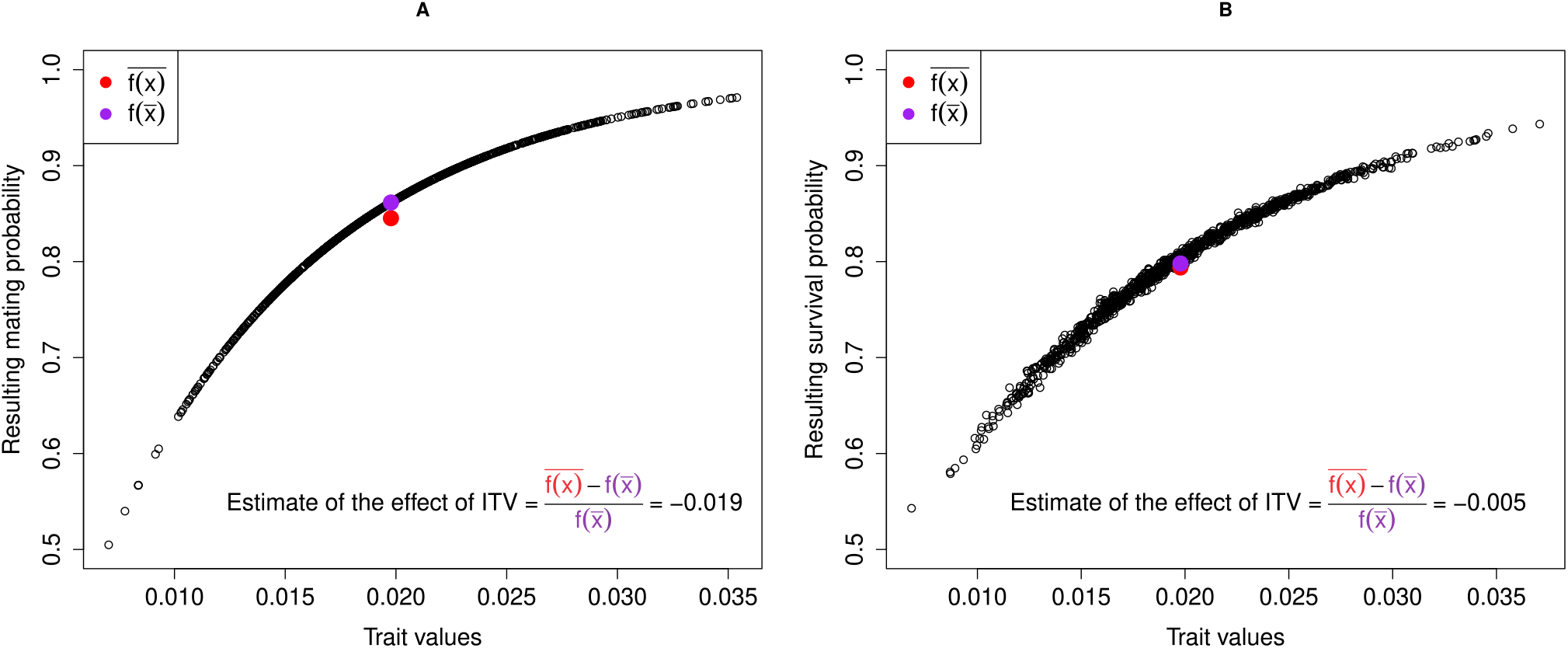
Estimates of the Jensen gap from 1000 randomly drawn trait values for (A) the mating probability in the mate-finding Allee effect model with 100 males present and (B) the survival probability in the predator-driven Allee effect model for a population size of 100. The red points depict the respective average probability of all used trait values while the purple points depict the probability corresponding to the average trait value, that is, to a population without ITV.

## Discussion

Here we have examined how population survival under two types of Allee effect, the mate-finding Allee effect and the predator-driven Allee effect, is impacted by intraspecific variation in relevant traits such as the mate-search rate and an anti-predator defense trait. Our analytical calculations and the simulation-based estimation of the Jensen gap showed that for both types of Allee effect, non-heritable intraspecific trait variation (ITV) reduced population growth (either through reduced mating probability or increased predation risk) and thus increased the Allee threshold. The results of the individual-based simulations closely matched with the analytical calculations. While the negative effect of non-heritable ITV was strong in the mate-finding Allee effect model, for the predator-driven Allee effect model it was too weak to be noticeable in the individual-based simulation results. Adding inheritance with a small mutational standard deviation mostly compensated the negative effects of ITV in the mate-finding Allee effect model, while larger mutational standard deviations further exacerbated the Allee effect. By contrast, for the predator-driven Allee effect model, inheritance and increasing mutational standard deviations had a positive effect on population survival.

The negative effects of ITV in both models arise via nonlinear averaging from the negative curvature of the mate-finding and survival functions. While the nonlinear averaging effect is straightforward in the mate-finding model because the mating probability only depends on the individual’s own trait, an individual’s survival probability under the predator-driven Allee effect also depends on the traits of the other individuals in the population. The reason for the weak effect of ITV in this model is that nonlinear averaging increases predation risk for focal individuals (the numerator of the expression), but also increases predation on other individuals in the population (via the denominator). Because the predator then spends more time handling other individuals, it has less time to hunt the focal individual, which buffers the effect of ITV in focal individuals. The net effect is still to increase predation and reduce survival probability, but the effect is weak.

Adding evolution, that is mutations and inheritance, has three effects with partly opposite consequences. First, there is input of additional variation via mutations, which tends to increase ITV and thus is expected to have a negative effect on population persistence. Second, there is selection of individuals with higher mate-search rates or better defense against predators, which should have a positive effect on survival. Third, inheritance can also lead to the evolution of a smaller trait variance if the mutation standard deviation is small, which again can have a positive effect on population survival. The third effect can explain why survival probabilities in the mate-finding Allee effect model with inheritance only or with ITV and inheritance and a small mutation standard deviation were approximately equal to those without ITV and inheritance. The level of ITV had equilibrated at a very low level, which then had hardly any effect anymore. For higher levels of the mutation standard deviation in the mate-finding Allee effect model, the first effect seems to dominate. That is, variance increases over time, which then reduces the mating probability and thus population persistence probability. By contrast, for the predator-driven Allee effect model, the second effect seems to dominate, at least for high enough mutation standard deviations. Here trait means increase substantially, thereby reducing predation pressure and increasing population persistence probabilities.

That evolution has clearer positive effects in the predator-driven Allee effect model than in the mate-finding Allee effect model can be explained as follows. In the mate-finding model, only the females search mates and make use of their trait value while the trait value of the males is irrelevant. Yet the trait values of the fathers still get inherited to their offspring, which leads to weaker overall selection pressures. Additionally, selection pressures in the mate-finding model lessen as the population gets larger and the mating probability approaches 1. In the predator-driven Allee effect model, by contrast, both sexes are subject to natural selection and selection stays constant over time and across densities. Finally, since the negative effects of ITV per se are weaker in the predator-driven Allee effect model it is easier for evolution to overpower these effects than in the mate-finding Allee effect model.

### Comparison with other studies

Among the different mechanisms mentioned by Bolnick et al. (2011) via which ITV can have an ecological effect, Jensen’s inequality (nonlinear averaging), phenotypic subsidy, eco-evolutionary dynamics, and trait sampling are relevant for our results. First, the negative effects of both higher trait and mutation standard deviation can be explained by Jensen’s inequality. Due to the curve of the function used to calculate the mating probability there is a negative effect of ITV. This means that overall it is better if all individuals share the same trait value instead of having them distributed around the mean either due to ITV or mutation. Our findings for the mate-finding Allee effect model are consistent with the observation of Berec (2018) that individual variation in mate-search rate has a negative effect on mating probability and help explain this observation in the light of nonlinear averaging. Second, the phenotypic subsidy effect in our model is that parents do not produce offspring that are identical to themselves, but new variance is generated in the offspring via mutation, which reduces fitness since variation has a detrimental effect in our models. Third, ITV in our model versions with inheritance can contribute to driving eco-evolutionary dynamics. As we have seen, evolutionary change can buffer or overpower the negative effects of ITV in some cases. Finally, the random sampling of traits in the initial population also contributes to the ITV effects in our case, even in the models with inheritance.

Although they did not focus on the effects of ITV, two previous studies (Berec et al., 2018; Watts and Fitzpatrick, 2026) have modeled the eco-evolutionary dynamics of mate-search rates. While Berec et al. (2018) used an individual-based simulation model where the mate-search rate is a polygenic trait with additive contributions across loci, Watts and Fitzpatrick (2026) used the adaptive dynamics approach to study the evolution of a trait that increases the encounter rate with mates. In contrast to our study, both studies assumed that increased mate-search rate can be costly. Watts and Fitzpatrick (2026) find that for each parameter combination, there is a single eco-evolutionary end-point with a certain evolutionarily stable mate-search rate and a certain equilibrium population size. However, since they focus on equilibria at which the population is present, they do not consider extinctions. By contrast, Berec et al. (2018) do allow for extinctions. Under some scenarios, mate-search rate in their model also evolves to an equilibrium value, but there are also situations where there is run-away evolution to higher and higher mate-search rates and then population extinction because of the high costs (evolutionary suicide). By contrast, we have assumed that the traits can evolve freely, without any costs. However, over the time scale considered, we did not observe unrealistically large changes in trait values. Thus our model is kind of a best-case scenario for the short-term benefit of evolution from the perspective of the modeled species. To make longer-term predictions, trade-offs and costs should be included.

### Model limitations and future perspectives

Although in both our models ITV reduced population survival probabilities and increased Allee thresholds, we cannot make a general conclusion that ITV worsens Allee effects. For example, for the predator-driven Allee effect, the negative effect was driven by the curvature of the assumed attack rate function. Individual variation in traits that have another functional relationship with fitness may lead to different results, as also evidenced by the negative effect of variation in search straightness on mating probability observed by Berec (2018). As a next step, one could for example explore ITV in prey traits like body width that can change the handling time of a predator (Hammill et al., 2015; Milonas et al., 2011). In each such case, it is thus important to find out the relationship between trait and predator attack rate. In cases where the relationship has different curvatures in different parts, the direction of the effect of ITV can even change for the same trait depending on the mean trait value (Pan et al., 2025; van Benthem et al., 2025).

In our model, individual traits were fixed and their expression did not depend on population density or other environmental variables. For future models, it could be interesting to also take phenotypic plasticity into account. For example, in the bush cricket *Roeseliana roeselii* (formerly *Metrioptera roeselii*) individuals covered larger distances when at low density and fertilization rate remained high (Kindvall et al., 1998). For such species, models that assume fixed traits might make overly pessimistic predictions for population survival at low density.

More generally, a more detailed understanding of the role of ITV in Allee effects will require more mechanistic models for specific systems. For example, one could create more complex spatial models that simulate actual movement during mate search (Berec, 2018; McCarthy, 1997) or keep track of the position of individuals in a “selfish herd” (Hamilton, 1971; Morton et al., 1994; Reluga and Viscido, 2005). One could also use a more sophisticated model in terms of the genetics and inheritance. This would make sense especially if the genetic basis of the trait of interest is known. Rare examples where this is the case are some morphological or chemical plant defense traits (Kawagoe et al., 2011; Olsen et al., 2013). However, these traits are then often not continuous but discrete so that it is harder to compare populations with the same mean trait and different levels of ITV.

Finally, it would be interesting to test the predicted effects of ITV for Allee effects with model organisms in the lab. For other questions, like for the role of ITV in species interactions, such experiments have been done (e.g. Hausch et al., 2018; Pan et al., 2025) by mixing different numbers of genetic lineages or by painting leaves with defense chemicals in a way to achieve a desired level of ITV. Such experiments would allow one to quantify mating probabilities or predation risk relatively easily, but to quantify Allee thresholds, many replicate populations with different founder population sizes would be necessary, which is often not feasible.

### Implications for conservation and invasion biology

In practice, these findings can be of varied use. For example, through the release of sex pheromones, the Allee threshold of an unwanted population with mate-finding Allee effect can be increased, leading to its extinction at higher population sizes (Berec et al., 2007; Fauvergue, 2013). Similarly releasing sterile males in an unwanted population can also interfere with a mate-finding Allee effect and increase the Allee threshold (Berec et al., 2007; Fauvergue, 2013). When releasing biological control agents to deal with a pest it is important to understand whether they suffer from a mate-finding Allee effect to ensure their survival (Fauvergue, 2013). We are not aware of any studies on the effect of ITV on these methods and it is likely hard to measure or even control it in wild or released populations. But this study shows that ITV impacts mate-finding Allee effects and therefore might be critical for the success of these population management strategies and should be looked at in the future.

Similar to the mate-finding Allee effect, the predator-driven Allee effect can be used in managing unwanted pest populations (Berec et al., 2007). Introduced predators or parasitoids can cause such an effect depending on their functional response (Berec et al., 2007). This study suggests that inheritance and ITV introduced through mutation affect the outcome of a predator-driven Allee effect. Understanding the impact ITV has on the interaction between pest and biological control agent could improve the process of managing pest populations. Furthermore, it is essential to assess a potential predator-driven Allee effect when managing threatened populations (Mooring et al., 2004). Also the outcome of translocations and reintroductions can be impacted by Allee effects (Mooring et al., 2004). Once again, the role of ITV in such measures is largely unknown but, if understood, could help improve their success.

## Supporting information

R code and simulation data

Maxima code for analytical apprimation

## Data and Code Availability

All R code used to generate the results and figures is provided as a supplementary zip file and will later be uploaded to a repository.

## Supplementary Information

**Figure S1:**
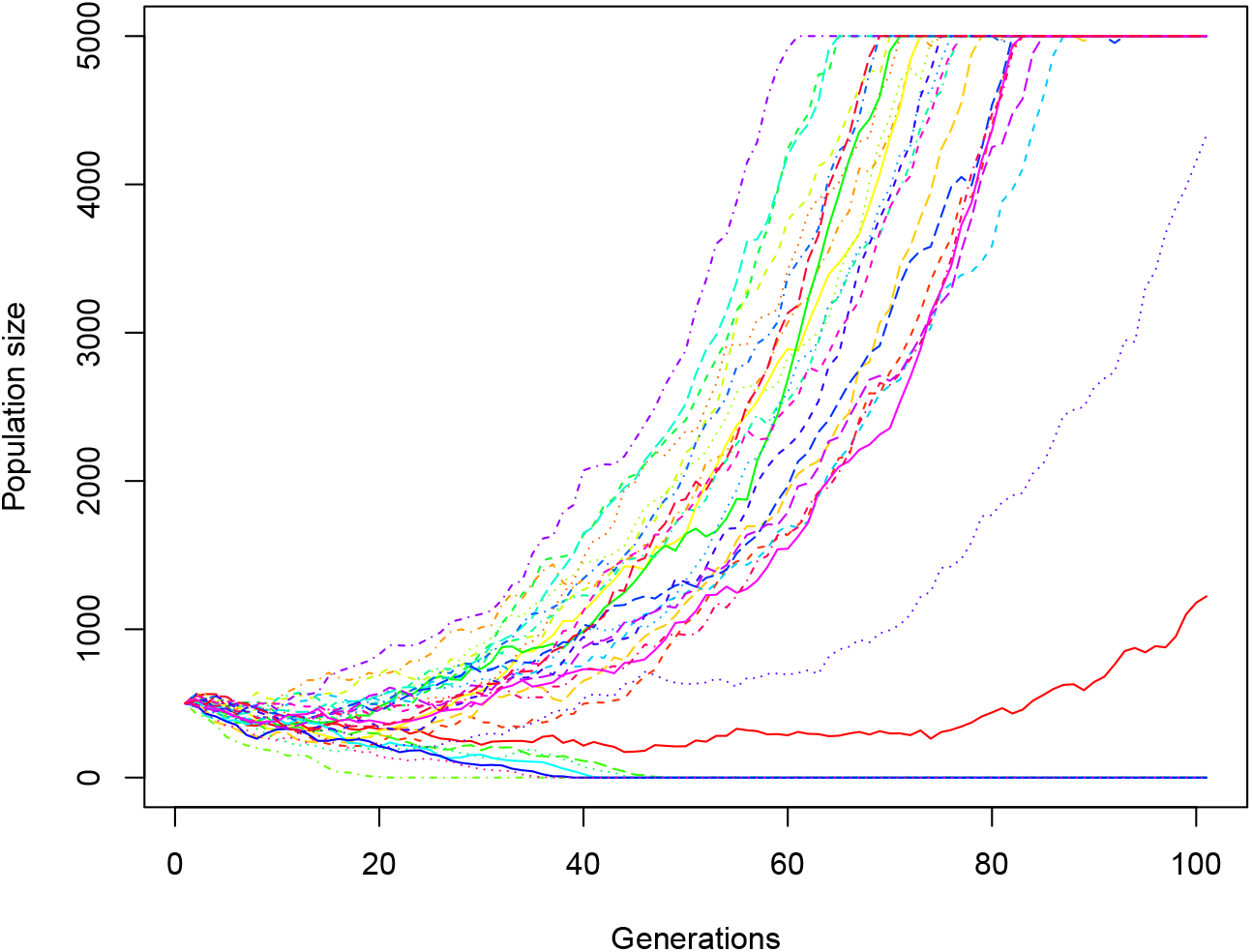
Example population-size trajectories for the first 30 individual replicates of the mate-finding Allee effect model with ITV and inheritance and a mutation standard deviation of 0.01, an initial population size of 500 and otherwise default parameters (corresponding to the dotted line in Fig. 2 A).

**Figure S2:**
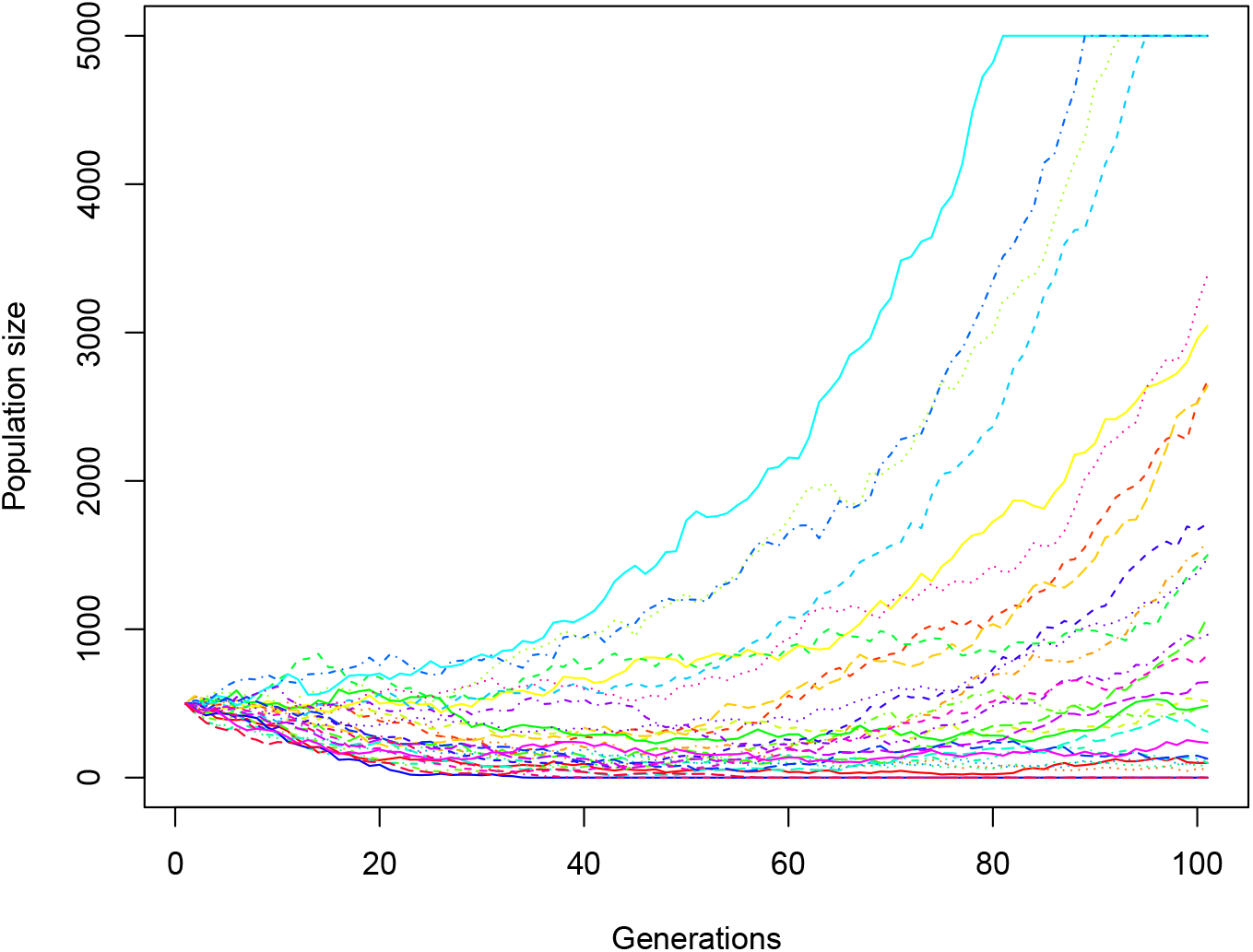
Example population-size trajectories for the first 30 individual replicates of the predator-driven Allee effect model with ITV and inheritance and a mutation standard deviation of 0.01, an initial population size of 500 and otherwise default parameters (corresponding to the dotted line in Fig. 4 A).

