## Supplementary material for "How does individual trait variation impact the survival of populations with an Allee effect?": R code and simulation data: All_functions_mate_finding.R

```
#Created 28.04.2025
####Standard function####

alleefunc <- function(years = 100, #Number of years
                      N0 = 500, #Starting population size
                      K = 5000, #Carrying capacity
                      percF = 0.5, #percentage (or better chance) of individual being female 
                      r = 0.02, #per capita encounter rate (female finding a male to mate with)
                      w = 2.1, #fecundity parameter
                      traitsd = 0.005, #Standard deviation of the trait r
                      mutation_sd = 0.0005, #Standard deviation of the mutation 
                      traitmin = 0, #Minimum value the trait can be (for the mutation function)
                      replicates = 100){ #number of replicates
  mating <- function(M,r){ #Function to calculate the mating probability of each individual
    1-exp(-r*M)
  }
  mutate <- function(old_trait,mutation_sd,trait_min){ #mutation function to change the value for r
    new_trait <- old_trait + rnorm(length(old_trait),0,mutation_sd)
    new_trait[which(new_trait<trait_min)] <- trait_min #prevent r from getting lower than 0
    return(new_trait)
  }
  
  repmatrix <- matrix(nrow = years+1,ncol = replicates) #Matrix to store pop size for each replicate
  meanrmatrix <- matrix(nrow = years+1,ncol = replicates) #Matrix to store mean r for each replicate
  sdrmatrix <- matrix(nrow = years+1,ncol = replicates) #Matrix to store the standard deviation of r for each replicate
  
  for (rep in 1:replicates) { #Start of the replicate loop
    
    popvec <- sample(c("F","M"),size=N0, replace = TRUE,prob = c(percF,1-percF)) #Generate initial population
    rvec <- rgamma(n = length(popvec), shape = (r/traitsd)^2 ,rate = r/traitsd^2) #Generate r value for each individual
    
    #Vectors to store each sex individually
    rvecF <- rvec[which(popvec=="F")] 
    rvecM <- rvec[which(popvec=="M")]
    
    #Vector to store population size over multiple time steps
    popsize <- numeric(years+1)
    popsize[1] <- N0
    
    #Vector to store the mean r
    meanr <- numeric(years+1)
    meanr[1] <- mean(rvec)
    sdrmatrix[1,rep] <- sd(rvec)
    
    for (t in 1:years) {
      N <- popsize[t] #current population size
      females <- sum(popvec=="F") #current number of females
      males <- sum(popvec=="M") #current number of males
      offspringvec <- numeric(females) #To prevent that a population survives with only males
      offspring_rvec <- numeric() #Empty vector to store r value of offspring
      
      if (females > 0){ 
        matingvec <- mating(males,rvecF) #Calculate the mating probability for each female
        mated <- rbinom(females,1,matingvec) #Calculating whether or not a female actually mates
        fathers <- sample(1:sum(popvec=="M"),sum(mated),replace = TRUE) #Drawing the mate for each female that mates; When false it could be that more females mate than mates are available
        offspringvec[mated==1] <- rpois(sum(mated),w) #Drawing number of offspring for each pair
        
        rvecMothers <- rvecF[which(mated==1)] #Get r value for each mated female
        rvecFathers <- rvecM[fathers] #Get r value for each father
        
        parentmatrix <- cbind(rvecMothers,rvecFathers) #Combine r value from both parents in a matrix
        
        meanrvec <- rowMeans(parentmatrix) #Take the mean value for r between the two parents
        
        offspring_rvec <- rep(meanrvec,offspringvec[which(mated==1)]) #Inherit the value for r to the offspring
        offspring_rvec <- mutate(offspring_rvec,mutation_sd,traitmin) #Mutate the offspring values for r
      }
      rvec <- offspring_rvec #Replace the current generation with the offspring
      
      if(length(rvec) > K) { #Reducing the population size if they are over the carrying cap
        over_cap <- sample(1:length(rvec),length(rvec)-K,replace = FALSE)
        rvec <- rvec[-over_cap]
        } 
      
      popsize[t+1] <- length(rvec) 
      
      meanr[t+1] <- mean(rvec) #Saving the mean r of the population
      sdrmatrix[t+1,rep] <- sd(rvec) #saving the standard deviation
      
      popvec <- sample(c("F","M"),size = popsize[t+1], replace = TRUE,prob = c(percF,1-percF)) #Generate sex for the next generation
      
      rvecF <- rvec[which(popvec=="F")] #Get r of all females 
      rvecM <- rvec[which(popvec=="M")] #Get r of all males
    }
    repmatrix[,rep] <- popsize #Save popsizevector of the replicate
    meanrmatrix[,rep] <- meanr #Save meanrvector of the replicate
  }
  survivalmatrix <- repmatrix[years+1,]>0 #Calculate if a population survived a simulation 
  return(list(repmatrix,meanrmatrix,survivalmatrix,sdrmatrix)) #Return results
}


###Function without inheritance####


noinheritance <- function(years = 100, #Number of years
                          N0 = 500, #Starting population size
                          K = 5000, #Carrying capacity
                          percF = 0.5, #percentage of individual being female 
                          r = 0.02, #per capita encounter rate (female finding a male to mate with)
                          w = 2.1, #fecundity parameter
                          traitsd = 0.005, #Standard deviation of the trait r
                          mutation_sd = 0.0005, #Standard deviation of the mutation 
                          traitmin = 0, #Minimum value the trait can be (for the mutation function)
                          replicates = 100){ #number of replicates
  mating <- function(M,r){ #Function to calculate the mating probability of each individual
    1-exp(-r*M)
  }
  #mutate <- function(old_trait,mutation_sd,trait_min){ #mutation function to change the value for r
  #new_trait <- old_trait + rnorm(length(old_trait),0,mutation_sd)
  #new_trait[which(new_trait<trait_min)] <- trait_min #prevent r from getting lower than 0
  #return(new_trait)
  #}
  
  repmatrix <- matrix(nrow = years+1,ncol = replicates) #Matrix to store pop size for each replicate
  meanrmatrix <- matrix(nrow = years+1,ncol = replicates) #Matrix to store mean r for each replicate
  sdrmatrix <- matrix(nrow = years+1,ncol = replicates) #Matrix to store the standard deviation of r for each replicate
  
  for (rep in 1:replicates) { #Start of the replicate loop
    
    popvec <- sample(c("F","M"),size=N0, replace = TRUE,prob = c(percF,1-percF)) #Generate initial population
    rvec <- rgamma(n = length(popvec), shape = (r/traitsd)^2 ,rate = r/traitsd^2) #Generate r value for each individual
    
    #Vectors to store each sex individually
    rvecF <- rvec[which(popvec=="F")]
    rvecM <- rvec[which(popvec=="M")]
    
    #Vector to store population size over multiple time steps
    popsize <- numeric(years+1)
    popsize[1] <- N0
    
    #Vector to store the mean r
    meanr <- numeric(years+1)
    meanr[1] <- mean(rvec)
    sdrmatrix[1,rep] <- sd(rvec)
    
    for (t in 1:years) {
      N <- popsize[t] #current population size
      females <- sum(popvec=="F") #current number of females
      males <- sum(popvec=="M") #current number of males
      offspringvec <- numeric(females) #To prevent that a population survives with only males
      offspring_rvec <- numeric() #Empty vector to store r value of offspring
      
      if (females > 0){
        matingvec <- mating(males,rvecF) #Calculate the mating probability for each female
        mated <- rbinom(females,1,matingvec) #Calculating whether or not a female actually mates
        #fathers <- sample(1:sum(popvec=="M"),sum(mated),replace = TRUE) #Drawing the mate for each female that mates; When false it could be that more females mate than mates are available
        offspringvec[mated==1] <- rpois(sum(mated),w) #Drawing number of offspring for each pair
        
        offspring_rvec <- rgamma(n = sum(offspringvec), shape = (r/traitsd)^2 ,rate = r/traitsd^2) #Draw rvec for offspring
        
        #rvecMothers <- rvecF[which(mated==1)] #Get r value for each mated female
        #rvecFathers <- rvecM[fathers] #Get r value for each father
        
        #parentmatrix <- cbind(rvecMothers,rvecFathers) #Combine r value from both parents in a matrix
        
        #meanrvec <- rowMeans(parentmatrix) #Take the mean value for r between the two parents
        
        #rvec <- rep(meanrvec,offspringvec[which(mated==1)]) #Inherit the value for r to the offspring
        #rvec <- mutate(rvec,mutation_sd,traitmin) #Mutate the offspring values for r
        
      }
      rvec <- offspring_rvec #Replace the current generation with the offspring
      
      if(length(rvec) > K) { #Reducing the population size if they are over the carrying cap
        over_cap <- sample(1:length(rvec),length(rvec)-K,replace = FALSE)
        rvec <- rvec[-over_cap]
      } 
      
      popsize[t+1] <- length(rvec) 
      meanr[t+1] <- mean(rvec) #Saving the mean r of the population
      sdrmatrix[t+1,rep] <- sd(rvec) #saving the standard deviation
      
      popvec <- sample(c("F","M"),size = popsize[t+1], replace = TRUE,prob = c(percF,1-percF)) #Generate sex for the next generation
      
      rvecF <- rvec[which(popvec=="F")] #Get r of all females 
      rvecM <- rvec[which(popvec=="M")] #Get r of all males
    }
    repmatrix[,rep] <- popsize #Save popsizevector of the replicate
    meanrmatrix[,rep] <- meanr #Save meanrvector of the replicate
  }
  survivalmatrix <- repmatrix[years+1,]>0 #Calculate if a population survived a simulation 
  return(list(repmatrix,meanrmatrix,survivalmatrix,sdrmatrix)) #Return results
} 


###Function without ITV####


noITV <- function(years = 100, #Number of years
                  N0 = 500, #Starting population size
                  K = 5000, #Carrying capacity
                  percF = 0.5, #percentage of individual being female 
                  r = 0.02, #per capita encounter rate (female finding a male to mate with)
                  w = 2.1, #fecundity parameter
                  traitsd = 0.005, #Standard deviation of the trait r
                  mutation_sd = 0.0005, #Standard deviation of the mutation 
                  traitmin = 0, #Minimum value the trait can be (for the mutation function)
                  replicates = 100){ #number of replicates
  mating <- function(M,r){ #Function to calculate the mating probability of each individual
    1-exp(-r*M)
  }
  mutate <- function(old_trait,mutation_sd,trait_min){ #mutation function to change the value for r
    new_trait <- old_trait + rnorm(length(old_trait),0,mutation_sd)
    new_trait[which(new_trait<trait_min)] <- trait_min #prevent r from getting lower than 0
    return(new_trait)
  }
  
  repmatrix <- matrix(nrow = years+1,ncol = replicates) #Matrix to store pop size for each replicate
  meanrmatrix <- matrix(nrow = years+1,ncol = replicates) #Matrix to store mean r for each replicate
  sdrmatrix <- matrix(nrow = years+1,ncol = replicates) #Matrix to store the standard deviation of r for each replicate
  
  for (rep in 1:replicates) { #Start of the replicate loop
    
    popvec <- sample(c("F","M"),size=N0, replace = TRUE,prob = c(percF,1-percF)) #Generate initial population
    #rvec <- rgamma(n = length(popvec), shape = (r/traitsd)^2 ,rate = r/traitsd^2) #Generate r value for each individual
    rvec <- rep(r,length(popvec))
    
    #Vectors to store each sex individually
    rvecF <- rvec[which(popvec=="F")]
    rvecM <- rvec[which(popvec=="M")]
    
    #Vector to store population size over multiple time steps
    popsize <- numeric(years+1)
    popsize[1] <- N0
    sdrmatrix[1,rep] <- sd(rvec)
    
    #Vector to store the mean r
    meanr <- numeric(years+1)
    meanr[1] <- mean(rvec)
    
    for (t in 1:years) {
      N <- popsize[t] #current population size
      females <- sum(popvec=="F") #current number of females
      males <- sum(popvec=="M") #current number of males
      offspringvec <- numeric(females) #To prevent that a population survives with only males
      offspring_rvec <- numeric() #Empty vector to store r value of offspring
      
      if (females > 0){
        matingvec <- mating(males,rvecF) #Calculate the mating probability for each female
        mated <- rbinom(females,1,matingvec) #Calculating whether or not a female actually mates
        fathers <- sample(1:sum(popvec=="M"),sum(mated),replace = TRUE) #Drawing the mate for each female that mates; When false it could be that more females mate than mates are available
        offspringvec[mated==1] <- rpois(sum(mated),w) #Drawing number of offspring for each pair
        
        rvecMothers <- rvecF[which(mated==1)] #Get r value for each mated female
        rvecFathers <- rvecM[fathers] #Get r value for each father
        
        parentmatrix <- cbind(rvecMothers,rvecFathers) #Combine r value from both parents in a matrix
        
        meanrvec <- rowMeans(parentmatrix) #Take the mean value for r between the two parents
        
        offspring_rvec <- rep(meanrvec,offspringvec[which(mated==1)]) #Inherit the value for r to the offspring
        offspring_rvec <- mutate(offspring_rvec,mutation_sd,traitmin) #Mutate the offspring values for r
      }
      rvec <- offspring_rvec #Replace the current generation with the offspring
      
      if(length(rvec) > K) { #Reducing the population size if they are over the carrying cap
        over_cap <- sample(1:length(rvec),length(rvec)-K,replace = FALSE)
        rvec <- rvec[-over_cap]
      } 
      
      popsize[t+1] <- length(rvec) 
      meanr[t+1] <- mean(rvec) #Saving the mean r of the population
      sdrmatrix[t+1,rep] <- sd(rvec) #saving the standard deviation
      
      popvec <- sample(c("F","M"),size = popsize[t+1], replace = TRUE,prob = c(percF,1-percF)) #Generate sex for the next generation
      
      rvecF <- rvec[which(popvec=="F")] #Get r of all females 
      rvecM <- rvec[which(popvec=="M")] #Get r of all males
    }
    repmatrix[,rep] <- popsize #Save popsizevector of the replicate
    meanrmatrix[,rep] <- meanr #Save meanrvector of the replicate
  }
  survivalmatrix <- repmatrix[years+1,]>0 #Calculate if a population survived a simulation 
  return(list(repmatrix,meanrmatrix,survivalmatrix,sdrmatrix)) #Return results
}


###Function without ITV and without inheritance####

noinheritance_ITV <- function(years = 100, #Number of years
                              N0 = 500, #Starting population size
                              K = 5000, #Carrying capacity
                              percF = 0.5, #percentage of individual being female 
                              r = 0.02, #per capita encounter rate (female finding a male to mate with)
                              w = 2.1, #fecundity parameter
                              traitsd = 0.005, #Standard deviation of the trait r
                              mutation_sd = 0.0005, #Standard deviation of the mutation 
                              traitmin = 0, #Minimum value the trait can be (for the mutation function)
                              replicates = 100){ #number of replicates
  mating <- function(M,r){ #Function to calculate the mating probability of each individual
    1-exp(-r*M)
  }
  #mutate <- function(old_trait,mutation_sd,trait_min){ #mutation function to change the value for r
  #new_trait <- old_trait + rnorm(length(old_trait),0,mutation_sd)
  #new_trait[which(new_trait<trait_min)] <- trait_min #prevent r from getting lower than 0
  #return(new_trait)
  #}
  
  repmatrix <- matrix(nrow = years+1,ncol = replicates) #Matrix to store pop size for each replicate
  meanrmatrix <- matrix(nrow = years+1,ncol = replicates) #Matrix to store mean r for each replicate
  sdrmatrix <- matrix(nrow = years+1,ncol = replicates) #Matrix to store the standard deviation of r for each replicate
  
  for (rep in 1:replicates) { #Start of the replicate loop
    
    popvec <- sample(c("F","M"),size=N0, replace = TRUE,prob = c(percF,1-percF)) #Generate initial population
    #rvec <- rgamma(n = length(popvec), shape = (r/traitsd)^2 ,rate = r/traitsd^2) #Generate r value for each individual
    
    #Vectors to store each sex individually
    rvec <- rep(r,length(popvec))
    
    rvecF <- rvec[which(popvec=="F")]
    rvecM <- rvec[which(popvec=="M")]
    
    #Vector to store population size over multiple time steps
    popsize <- numeric(years+1)
    popsize[1] <- N0
    
    #Vector to store the mean r
    meanr <- numeric(years+1)
    meanr[1] <- mean(rvec)
    sdrmatrix[1,rep] <- sd(rvec)
    
    for (t in 1:years) {
      N <- popsize[t] #current population size
      females <- sum(popvec=="F") #current number of females
      males <- sum(popvec=="M") #current number of males
      offspringvec <- numeric(females) #To prevent that a population survives with only males
      offspring_rvec <- numeric() #Empty vector to store r value of offspring
      
      if (females > 0){
        matingvec <- mating(males,rvecF) #Calculate the mating probability for each female
        mated <- rbinom(females,1,matingvec) #Calculating whether or not a female actually mates
        #fathers <- sample(1:sum(popvec=="M"),sum(mated),replace = TRUE) #Drawing the mate for each female that mates; When false it could be that more females mate than mates are available
        offspringvec[mated==1] <- rpois(sum(mated),w) #Drawing number of offspring for each pair
        
        offspring_rvec <- rep(r,sum(offspringvec))  #Draw rvec for offspring (use sum since all have the same value)
        
        #rvecMothers <- rvecF[which(mated==1)] #Get r value for each mated female
        #rvecFathers <- rvecM[fathers] #Get r value for each father
        
        #parentmatrix <- cbind(rvecMothers,rvecFathers) #Combine r value from both parents in a matrix
        
        #meanrvec <- rowMeans(parentmatrix) #Take the mean value for r between the two parents
        
        #rvec <- rep(meanrvec,offspringvec[which(mated==1)]) #Inherit the value for r to the offspring
        #rvec <- mutate(rvec,mutation_sd,traitmin)  #Mutate the offspring values for r
        
      }
      rvec <- offspring_rvec #Replace the current generation with the offspring
      
      if(length(rvec) > K) { #Reducing the population size if they are over the carrying cap
        over_cap <- sample(1:length(rvec),length(rvec)-K,replace = FALSE)
        rvec <- rvec[-over_cap]
      } 
      
      popsize[t+1] <- length(rvec) 
      meanr[t+1] <- mean(rvec) #Saving the mean r of the population
      sdrmatrix[t+1,rep] <- sd(rvec) #saving the standard deviation
      
      popvec <- sample(c("F","M"),size = popsize[t+1], replace = TRUE,prob = c(percF,1-percF)) #Generate sex for the next generation
      
      rvecF <- rvec[which(popvec=="F")] #Get r of all females 
      rvecM <- rvec[which(popvec=="M")] #Get r of all males
    }
    repmatrix[,rep] <- popsize #Save popsizevector of the replicate
    meanrmatrix[,rep] <- meanr #Save meanrvector of the replicate
  }
  survivalmatrix <- repmatrix[years+1,]>0 #Calculate if a population survived a simulation 
  return(list(repmatrix,meanrmatrix,survivalmatrix, sdrmatrix)) #Return results
} 
```
