## Supplementary material for "How does individual trait variation impact the survival of populations with an Allee effect?": R code and simulation data: All_functions_predator_prey.R

```
#Created 28.04.2025
####Standard function####

alleefunc_pred <- function(years = 100, #Number of years
                      N0 = 500, #Starting population size
                      K = 5000, #Carrying capacity
                      percF = 0.5, #percentage (or better chance) of individual being female 
                      trait = 0.02, #individual trait value determining the decline of the attack rate
                      P = 1, #Number of Predators
                      c = 2, # attack rate of the predator on the smallest possible trait value (max attack rate)
                      b = 70, #decay of the attack rate with increasing trait value
                      h = 0.03, #handling time of the predator for one prey individual
                      w = 2.1, #fecundity parameter
                      traitsd = 0.005, #Standard deviation of the trait r
                      mutation_sd = 0.0005, #Standard deviation of the mutation 
                      traitmin = 0, #Minimum value the trait can be (for the mutation function)
                      replicates = 100){ #number of replicates
  survival_prob <- function(P,c,b,t,h,trait_vec){ #Function to calculate the survival probability of each individual
    1-((c*exp(-b*t))/(1+sum(c*exp(-b*trait_vec)*h))*P)
  }
  mutate <- function(old_trait,mutation_sd,trait_min){ #mutation function to change the value for r
    new_trait <- old_trait + rnorm(length(old_trait),0,mutation_sd)
    new_trait[which(new_trait<trait_min)] <- trait_min #prevent r from getting lower than 0
    return(new_trait)
  }
  
  repmatrix <- matrix(nrow = years+1,ncol = replicates) #Matrix to store pop size for each replicate
  meanrmatrix <- matrix(nrow = years+1,ncol = replicates) #Matrix to store mean r for each replicate
  sdrmatrix <- matrix(nrow = years+1,ncol = replicates) #Matrix to store the standard deviation of r for each replicate
  
  for (rep in 1:replicates) { #Start of the replicate loop
    
    popvec <- sample(c("F","M"),size=N0, replace = TRUE,prob = c(percF,1-percF)) #Generate initial population
    tvec <- rgamma(n = length(popvec), shape = (trait/traitsd)^2 ,rate = trait/traitsd^2) #Generate r value for each individual
    
    #Vectors to store each sex individually
    tvecF <- tvec[which(popvec=="F")] 
    tvecM <- tvec[which(popvec=="M")]
    
    #Vector to store population size over multiple time steps
    popsize <- numeric(years+1)
    popsize[1] <- N0
    
    #Vector to store the mean r
    meanr <- numeric(years+1)
    meanr[1] <- mean(tvec)
    sdrmatrix[1,rep] <- sd(tvec)
    
    for (t in 1:years) {
      N <- popsize[t] #current population size
      females <- sum(popvec=="F") #current number of females
      males <- sum(popvec=="M") #current number of males
      offspringvec <- numeric(females) #To prevent that a population survives with only males
      offspring_tvec <- numeric() #Empty vector to store r value of offspring
      survivalvec_f <- 0
      survivalvec_m <- 0
      
      if (females > 0){ 
        survivalvec_f <- survival_prob(P = P,c = c,b = b,t = tvecF,h = h,trait_vec = c(tvecF,tvecM)) #Calculate the survival probability of each female 
        survivalvec_m <- survival_prob(P = P,c = c,b = b,t = tvecM,h = h,trait_vec = c(tvecF,tvecM)) #Calculate the survival probability of each male 
        
        survivalvec_f[which(survivalvec_f<0)] <- 0 #Replacing any values below zero with 0
        survivalvec_m[which(survivalvec_m<0)] <- 0 #Replacing any values below zero with 0   
        
        survived_f <- rbinom(females,1, survivalvec_f) #Calculating whether or not a female actually survives
        survived_m <- rbinom(males,1, survivalvec_m) #Calculating whether or not a male actually survives
        tvecF <- tvecF[which(survived_f == 1)] #Removing dead females
        tvecM <- tvecM[which(survived_m == 1)] #Removing dead males
      if (length(tvecF) > 0 && length(tvecM) > 0){  
        fathers <- sample(1:length(tvecM),length(tvecF),replace = TRUE) #Drawing the mate for each female that survives; When false it could be that more females mate than mates are available
        offspringvec[survived_f == 1] <- rpois(sum(survived_f),w) #Drawing number of offspring for each pair
        
        tvecMothers <- tvecF #Get r value for each surviving female (all reproduce)
        tvecFathers <- tvecM[fathers] #Get r value for each father
        
        parentmatrix <- cbind(tvecMothers,tvecFathers) #Combine r value from both parents in a matrix
        
        meantvec <- rowMeans(parentmatrix) #Take the mean value for r between the two parents
        
        offspring_tvec <- rep(meantvec,offspringvec[which(survived_f == 1)]) #Inherit the value for r to the offspring
        offspring_tvec <- mutate(offspring_tvec,mutation_sd,traitmin) #Mutate the offspring values for r
       }
      }
      tvec <- offspring_tvec #Replace the current generation with the offspring
      
      if(length(tvec) > K) { #Reducing the population size if they are over the carrying cap
        over_cap <- sample(1:length(tvec),length(tvec)-K,replace = FALSE)
        tvec <- tvec[-over_cap]
      } 
      
      popsize[t+1] <- length(tvec) 
      meanr[t+1] <- mean(tvec) #Saving the mean r of the population
      sdrmatrix[t+1,rep] <- sd(tvec) #saving the standard deviation
      
      popvec <- sample(c("F","M"),size = popsize[t+1], replace = TRUE,prob = c(percF,1-percF)) #Generate sex for the next generation
      
      tvecF <- tvec[which(popvec=="F")] #Get r of all females 
      tvecM <- tvec[which(popvec=="M")] #Get r of all males
      
    }
    repmatrix[,rep] <- popsize #Save popsizevector of the replicate
    meanrmatrix[,rep] <- meanr #Save meantvector of the replicate
  }
  survivalmatrix <- repmatrix[years+1,]>0 #Calculate if a population survived a simulation 
  return(list(repmatrix,meanrmatrix,survivalmatrix,sdrmatrix)) #Return results
}


###Function without inheritance####


noinheritance_pred <- function(years = 100, #Number of years
                          N0 = 500, #Starting population size
                          K = 5000, #Carrying capacity
                          percF = 0.5, #percentage (or better chance) of individual being female 
                          trait = 0.02, #individual trait value determining the decline of the attack rate
                          P = 1, #Number of Predators
                          c = 2, # attack rate of the predator on the smallest possible trait value (max attack rate)
                          b = 70, #decay of the attack rate with increasing trait value
                          h = 0.03, #handling time of the predator for one prey individual
                          w = 2.1, #fecundity parameter
                          traitsd = 0.005, #Standard deviation of the trait r
                          mutation_sd = 0.0005, #Standard deviation of the mutation 
                          traitmin = 0, #Minimum value the trait can be (for the mutation function)
                          replicates = 100){ #number of replicates
  survival_prob <- function(P,c,b,t,h,trait_vec){ #Function to calculate the survival probability of each individual
    1-((c*exp(-b*t))/(1+sum(c*exp(-b*trait_vec)*h))*P)
  }
  #mutate <- function(old_trait,mutation_sd,trait_min){ #mutation function to change the value for r
  #new_trait <- old_trait + rnorm(length(old_trait),0,mutation_sd)
  #new_trait[which(new_trait<trait_min)] <- trait_min #prevent r from getting lower than 0
  #return(new_trait)
  #}
  
  repmatrix <- matrix(nrow = years+1,ncol = replicates) #Matrix to store pop size for each replicate
  meanrmatrix <- matrix(nrow = years+1,ncol = replicates) #Matrix to store mean r for each replicate
  sdrmatrix <- matrix(nrow = years+1,ncol = replicates) #Matrix to store the standard deviation of r for each replicate
  
  for (rep in 1:replicates) { #Start of the replicate loop
    
    popvec <- sample(c("F","M"),size=N0, replace = TRUE,prob = c(percF,1-percF)) #Generate initial population
    tvec <- rgamma(n = length(popvec), shape = (trait/traitsd)^2 ,rate = trait/traitsd^2) #Generate r value for each individual
    
    #Vectors to store each sex individually
    tvecF <- tvec[which(popvec=="F")]
    tvecM <- tvec[which(popvec=="M")]
    
    #Vector to store population size over multiple time steps
    popsize <- numeric(years+1)
    popsize[1] <- N0
    
    #Vector to store the mean r
    meanr <- numeric(years+1)
    meanr[1] <- mean(tvec)
    sdrmatrix[1,rep] <- sd(tvec)
    
    for (t in 1:years) {
      N <- popsize[t] #current population size
      females <- sum(popvec=="F") #current number of females
      males <- sum(popvec=="M") #current number of males
      offspringvec <- numeric(females) #To prevent that a population survives with only males
      offspring_tvec <- numeric() #Empty vector to store r value of offspring
      survivalvec_f <- 0
      survivalvec_m <- 0
      
      if (females > 0){ 
        survivalvec_f <- survival_prob(P = P,c = c,b = b,t = tvecF,h = h,trait_vec = c(tvecF,tvecM)) #Calculate the survival probability of each female 
        survivalvec_m <- survival_prob(P = P,c = c,b = b,t = tvecM,h = h,trait_vec = c(tvecF,tvecM)) #Calculate the survival probability of each male 
        
        survivalvec_f[which(survivalvec_f<0)] <- 0 #Replacing any values below zero with 0
        survivalvec_m[which(survivalvec_m<0)] <- 0 #Replacing any values below zero with 0   
        
        survived_f <- rbinom(females,1, survivalvec_f) #Calculating whether or not a female actually survives
        survived_m <- rbinom(males,1, survivalvec_m) #Calculating whether or not a male actually survives
        tvecF <- tvecF[which(survived_f == 1)] #Removing dead females
        tvecM <- tvecM[which(survived_m == 1)] #Removing dead males
      if (length(tvecF) > 0 && length(tvecM) > 0){  
        fathers <- sample(1:length(tvecM),length(tvecF),replace = TRUE) #Drawing the mate for each female that survives; When false it could be that more females mate than mates are available
        offspringvec[survived_f == 1] <- rpois(sum(survived_f),w) #Drawing number of offspring for each pair
        
        #tvecMothers <- tvecF #Get r value for each surviving female (all reproduce)
        #tvecFathers <- tvecM[fathers] #Get r value for each father
        
        #parentmatrix <- cbind(tvecMothers,tvecFathers) #Combine r value from both parents in a matrix
        
        #meantvec <- rowMeans(parentmatrix) #Take the mean value for r between the two parents
        
        offspring_tvec <- tvec <- rgamma(n = sum(offspringvec), shape = (trait/traitsd)^2 ,rate = trait/traitsd^2) #Inherit the value for r to the offspring
        #offspring_tvec <- mutate(offspring_tvec,mutation_sd,traitmin) #Mutate the offspring values for r
       }
      }
      tvec <- offspring_tvec #Replace the current generation with the offspring
      
      if(length(tvec) > K) { #Reducing the population size if they are over the carrying cap
        over_cap <- sample(1:length(tvec),length(tvec)-K,replace = FALSE)
        tvec <- tvec[-over_cap]
      } 
      
      popsize[t+1] <- length(tvec) 
      meanr[t+1] <- mean(tvec) #Saving the mean r of the population
      sdrmatrix[t+1,rep] <- sd(tvec) #saving the standard deviation
      
      popvec <- sample(c("F","M"),size = popsize[t+1], replace = TRUE,prob = c(percF,1-percF)) #Generate sex for the next generation
      
      tvecF <- tvec[which(popvec=="F")] #Get r of all females 
      tvecM <- tvec[which(popvec=="M")] #Get r of all males
    }
    repmatrix[,rep] <- popsize #Save popsizevector of the replicate
    meanrmatrix[,rep] <- meanr #Save meantvector of the replicate
  }
  survivalmatrix <- repmatrix[years+1,]>0 #Calculate if a population survived a simulation 
  return(list(repmatrix,meanrmatrix,survivalmatrix,sdrmatrix)) #Return results
} 


###Function without ITV####


noITV_pred <- function(years = 100, #Number of years
                  N0 = 500, #Starting population size
                  K = 5000, #Carrying capacity
                  percF = 0.5, #percentage (or better chance) of individual being female 
                  trait = 0.02, #individual trait value determining the decline of the attack rate
                  P = 1, #Number of Predators
                  c = 2, # attack rate of the predator on the smallest possible trait value (max attack rate)
                  b = 70, #decay of the attack rate with increasing trait value
                  h = 0.03, #handling time of the predator for one prey individual
                  w = 2.1, #fecundity parameter
                  traitsd = 0.005, #Standard deviation of the trait r
                  mutation_sd = 0.0005, #Standard deviation of the mutation 
                  traitmin = 0, #Minimum value the trait can be (for the mutation function)
                  replicates = 100){ #number of replicates
  survival_prob <- function(P,c,b,t,h,trait_vec){ #Function to calculate the survival probability of each individual
    1-((c*exp(-b*t))/(1+sum(c*exp(-b*trait_vec)*h))*P)
  }
  mutate <- function(old_trait,mutation_sd,trait_min){ #mutation function to change the value for r
    new_trait <- old_trait + rnorm(length(old_trait),0,mutation_sd)
    new_trait[which(new_trait<trait_min)] <- trait_min #prevent r from getting lower than 0
    return(new_trait)
  }
  
  repmatrix <- matrix(nrow = years+1,ncol = replicates) #Matrix to store pop size for each replicate
  meanrmatrix <- matrix(nrow = years+1,ncol = replicates) #Matrix to store mean r for each replicate
  sdrmatrix <- matrix(nrow = years+1,ncol = replicates) #Matrix to store the standard deviation of r for each replicate
  
  for (rep in 1:replicates) { #Start of the replicate loop
    
    popvec <- sample(c("F","M"),size=N0, replace = TRUE,prob = c(percF,1-percF)) #Generate initial population
    #tvec <- rgamma(n = length(popvec), shape = (r/traitsd)^2 ,rate = r/traitsd^2) #Generate r value for each individual
    tvec <- rep(trait,length(popvec))
    
    #Vectors to store each sex individually
    tvecF <- tvec[which(popvec=="F")]
    tvecM <- tvec[which(popvec=="M")]
    
    #Vector to store population size over multiple time steps
    popsize <- numeric(years+1)
    popsize[1] <- N0
    
    #Vector to store the mean r
    meanr <- numeric(years+1)
    meanr[1] <- mean(tvec)
    sdrmatrix[1,rep] <- sd(tvec)
    
    for (t in 1:years) {
      N <- popsize[t] #current population size
      females <- sum(popvec=="F") #current number of females
      males <- sum(popvec=="M") #current number of males
      offspringvec <- numeric(females) #To prevent that a population survives with only males
      offspring_tvec <- numeric() #Empty vector to store r value of offspring
      survivalvec_f <- 0
      survivalvec_m <- 0
      
      if (females > 0){ 
        survivalvec_f <- survival_prob(P = P,c = c,b = b,t = tvecF,h = h,trait_vec = c(tvecF,tvecM)) #Calculate the survival probability of each female 
        survivalvec_m <- survival_prob(P = P,c = c,b = b,t = tvecM,h = h,trait_vec = c(tvecF,tvecM)) #Calculate the survival probability of each male 
        
        survivalvec_f[which(survivalvec_f<0)] <- 0 #Replacing any values below zero with 0
        survivalvec_m[which(survivalvec_m<0)] <- 0 #Replacing any values below zero with 0   
        
        survived_f <- rbinom(females,1, survivalvec_f) #Calculating whether or not a female actually survives
        survived_m <- rbinom(males,1, survivalvec_m) #Calculating whether or not a male actually survives
        tvecF <- tvecF[which(survived_f == 1)] #Removing dead females
        tvecM <- tvecM[which(survived_m == 1)] #Removing dead males
        if (length(tvecF) > 0 && length(tvecM) > 0){  
          fathers <- sample(1:length(tvecM),length(tvecF),replace = TRUE) #Drawing the mate for each female that survives; When false it could be that more females mate than mates are available
          offspringvec[survived_f == 1] <- rpois(sum(survived_f),w) #Drawing number of offspring for each pair
          
          tvecMothers <- tvecF #Get r value for each surviving female (all reproduce)
          tvecFathers <- tvecM[fathers] #Get r value for each father
          
          parentmatrix <- cbind(tvecMothers,tvecFathers) #Combine r value from both parents in a matrix
          
          meantvec <- rowMeans(parentmatrix) #Take the mean value for r between the two parents
          
          offspring_tvec <- rep(meantvec,offspringvec[which(survived_f == 1)]) #Inherit the value for r to the offspring
          offspring_tvec <- mutate(offspring_tvec,mutation_sd,traitmin) #Mutate the offspring values for r
        }
      }
      tvec <- offspring_tvec #Replace the current generation with the offspring
      
      if(length(tvec) > K) { #Reducing the population size if they are over the carrying cap
        over_cap <- sample(1:length(tvec),length(tvec)-K,replace = FALSE)
        tvec <- tvec[-over_cap]
      } 
      
      popsize[t+1] <- length(tvec) 
      meanr[t+1] <- mean(tvec) #Saving the mean r of the population
      sdrmatrix[t+1,rep] <- sd(tvec) #saving the standard deviation
      
      popvec <- sample(c("F","M"),size = popsize[t+1], replace = TRUE,prob = c(percF,1-percF)) #Generate sex for the next generation
      
      tvecF <- tvec[which(popvec=="F")] #Get r of all females 
      tvecM <- tvec[which(popvec=="M")] #Get r of all males
    }
    repmatrix[,rep] <- popsize #Save popsizevector of the replicate
    meanrmatrix[,rep] <- meanr #Save meantvector of the replicate
  }
  survivalmatrix <- repmatrix[years+1,]>0 #Calculate if a population survived a simulation 
  return(list(repmatrix,meanrmatrix,survivalmatrix,sdrmatrix)) #Return results
}


###Function without ITV and without inheritance####

noinheritance_ITV_pred <- function(years = 100, #Number of years
                              N0 = 500, #Starting population size
                              K = 5000, #Carrying capacity
                              percF = 0.5, #percentage (or better chance) of individual being female 
                              trait = 0.02, #individual trait value determining the decline of the attack rate
                              P = 1, #Number of Predators
                              c = 2, # attack rate of the predator on the smallest possible trait value (max attack rate)
                              b = 70, #decay of the attack rate with increasing trait value
                              h = 0.03, #handling time of the predator for one prey individual
                              w = 2.1, #fecundity parameter
                              traitsd = 0.005, #Standard deviation of the trait r
                              mutation_sd = 0.0005, #Standard deviation of the mutation 
                              traitmin = 0, #Minimum value the trait can be (for the mutation function)
                              replicates = 100){ #number of replicates
  survival_prob <- function(P,c,b,t,h,trait_vec){ #Function to calculate the survival probability of each individual
    1-((c*exp(-b*t))/(1+sum(c*exp(-b*trait_vec)*h))*P)
  }
  #mutate <- function(old_trait,mutation_sd,trait_min){ #mutation function to change the value for r
  #new_trait <- old_trait + rnorm(length(old_trait),0,mutation_sd)
  #new_trait[which(new_trait<trait_min)] <- trait_min #prevent r from getting lower than 0
  #return(new_trait)
  #}
  
  repmatrix <- matrix(nrow = years+1,ncol = replicates) #Matrix to store pop size for each replicate
  meanrmatrix <- matrix(nrow = years+1,ncol = replicates) #Matrix to store mean r for each replicate
  sdrmatrix <- matrix(nrow = years+1,ncol = replicates) #Matrix to store the standard deviation of r for each replicate
  
  for (rep in 1:replicates) { #Start of the replicate loop
    
    popvec <- sample(c("F","M"),size=N0, replace = TRUE,prob = c(percF,1-percF)) #Generate initial population
    #tvec <- rgamma(n = length(popvec), shape = (r/traitsd)^2 ,rate = r/traitsd^2) #Generate r value for each individual
    
    #Vectors to store each sex individually
    tvec <- rep(trait,length(popvec))
    
    tvecF <- tvec[which(popvec=="F")]
    tvecM <- tvec[which(popvec=="M")]
    
    #Vector to store population size over multiple time steps
    popsize <- numeric(years+1)
    popsize[1] <- N0
    
    #Vector to store the mean r
    meanr <- numeric(years+1)
    meanr[1] <- mean(tvec)
    sdrmatrix[1,rep] <- sd(tvec)
    
    for (t in 1:years) {
      N <- popsize[t] #current population size
      females <- sum(popvec=="F") #current number of females
      males <- sum(popvec=="M") #current number of males
      offspringvec <- numeric(females) #To prevent that a population survives with only males
      offspring_tvec <- numeric() #Empty vector to store r value of offspring
      survivalvec_f <- 0
      survivalvec_m <- 0
      
      if (females > 0){ 
        survivalvec_f <- survival_prob(P = P,c = c,b = b,t = tvecF,h = h,trait_vec = c(tvecF,tvecM)) #Calculate the survival probability of each female 
        survivalvec_m <- survival_prob(P = P,c = c,b = b,t = tvecM,h = h,trait_vec = c(tvecF,tvecM)) #Calculate the survival probability of each male 
        
        survivalvec_f[which(survivalvec_f<0)] <- 0 #Replacing any values below zero with 0
        survivalvec_m[which(survivalvec_m<0)] <- 0 #Replacing any values below zero with 0   
        
        survived_f <- rbinom(females,1, survivalvec_f) #Calculating whether or not a female actually survives
        survived_m <- rbinom(males,1, survivalvec_m) #Calculating whether or not a male actually survives
        tvecF <- tvecF[which(survived_f == 1)] #Removing dead females
        tvecM <- tvecM[which(survived_m == 1)] #Removing dead males
      if (length(tvecF) > 0 && length(tvecM) > 0){  
        fathers <- sample(1:length(tvecM),length(tvecF),replace = TRUE) #Drawing the mate for each female that survives; When false it could be that more females mate than mates are available
        offspringvec[survived_f == 1] <- rpois(sum(survived_f),w) #Drawing number of offspring for each pair
        
        #tvecMothers <- tvecF #Get r value for each surviving female (all reproduce)
        #tvecFathers <- tvecM[fathers] #Get r value for each father
        
        #parentmatrix <- cbind(tvecMothers,tvecFathers) #Combine r value from both parents in a matrix
        
        #meantvec <- rowMeans(parentmatrix) #Take the mean value for r between the two parents
        
        offspring_tvec <- rep(trait,sum(offspringvec))
        #offspring_tvec <- mutate(offspring_tvec,mutation_sd,traitmin) #Mutate the offspring values for r
       }
      }
      tvec <- offspring_tvec #Replace the current generation with the offspring
      
      if(length(tvec) > K) { #Reducing the population size if they are over the carrying cap
        over_cap <- sample(1:length(tvec),length(tvec)-K,replace = FALSE)
        tvec <- tvec[-over_cap]
      } 
      
      popsize[t+1] <- length(tvec) 
      meanr[t+1] <- mean(tvec) #Saving the mean r of the population
      sdrmatrix[t+1,rep] <- sd(tvec) #saving the standard deviation
      
      popvec <- sample(c("F","M"),size = popsize[t+1], replace = TRUE,prob = c(percF,1-percF)) #Generate sex for the next generation
      
      tvecF <- tvec[which(popvec=="F")] #Get r of all females 
      tvecM <- tvec[which(popvec=="M")] #Get r of all males
    }
    repmatrix[,rep] <- popsize #Save popsizevector of the replicate
    meanrmatrix[,rep] <- meanr #Save meantvector of the replicate
  }
  survivalmatrix <- repmatrix[years+1,]>0 #Calculate if a population survived a simulation 
  return(list(repmatrix,meanrmatrix,survivalmatrix,sdrmatrix)) #Return results
} 
```
