## Supplementary material for "How does individual trait variation impact the survival of populations with an Allee effect?": R code and simulation data: Fig1_Fig3.R

```
####Making the plot for mate finding Allee effect

load("Simulation_results/Mate_finding/Mate_finding_different_mutation_sd_500_replicates_0.001_0_005_0.01_pop_sequence_10_800_10_Schritte.RData")

#For some reason this dataset loads with a different name
load("Simulation_results/Mate_finding/Mate_finding_different_trait_sd_500_replicates_0.001_0_005_0.01_pop_sequence_10_800_10_Schritte.RData")

load("Simulation_results/Mate_finding/mate_finding_standard_all_population_sizes.RData")

#Renaming for easier use

mate_finding_standard <- mate_finding_standard_all_population_sizes

mate_finding_mutation_sd <- Mate_finding_different_mutation_sd_500_replicates_0.001_0_005_0.01_pop_sequence_10_800_10_Schritte

mate_finding_trait_sd <- Mate_finding_different_trait_sd_500_replicates_0.001_0_005_0.01_pop_sequence_10_800_10_Schritte

#Equation for the equilibrium population size

equilibriumpopulationsizeGamma<- function(mu,sigma,w){ 
  Nstar <- (2*mu*((w/(w-2))^(sigma^2/mu^2)-1))/sigma^2
  return(Nstar)
}

#Equation for the equilibrium population size if sigma is 0

equilibriumpopulationsizeGamma_None<- function(mu,w){ 
  Nstar <- 2/mu * log(w/(w-2))
  return(Nstar)
}

#Setting standard parameters used in the model for calculating the equilibrium population size

mu = 0.02 
sigma = 0.005
w = 2.1

#Calculating equilibrium population size 

AT_standard <- equilibriumpopulationsizeGamma(mu = mu, sigma = sigma, w = w)

#Calculating equilibrium population size if sigma is 0

AT_none <- equilibriumpopulationsizeGamma_None(mu = mu, w = w)

#Making the mate finding plots

par(mfrow=c(1,3))
par(mar = c(5.1, 4.5, 4.1, 0.1))

startingpop <- seq(10,800,1) #Starting population sizes

testvec <-c(0.001,0.005,0.01) #Values used when testing different values for the trait and mutation sd 

#Plot for the standard values

matplot(startingpop, mate_finding_standard[[1]],type = "l", xlab = "Founder population size", ylab = "Proportion of surviving populations",col = "#011C40FF", cex.axis = 1.8, cex.lab = 1.8, lwd = 3 , ylim = c(0,1), main = "A", cex.main = 1.8)
matlines(startingpop, mate_finding_standard[[2]],col = "#40e078", lwd = 3)
matlines(startingpop, mate_finding_standard[[3]], col = "#05DBF2FF", lwd = 3)
matlines(startingpop, mate_finding_standard[[4]], col = "#4C4CFFFF", lwd = 3)
abline(v = AT_standard, col = "#40e078", lwd = 3)
abline(v = AT_none, col = "#4C4CFFFF", lwd = 3)
legend("bottomright", title = "Model includes:", legend = c("ITV + Inheritance", "ITV only", "Inheritance only", "None"), col = c("#011C40FF", "#40e078", "#05DBF2FF", "#4C4CFFFF"), lty = rep(1,5), lwd = c(rep(2,4)) ,ncol = 1, cex = 1.3 , bty="o")

startingpop <- seq(10,800,10) #For these graphs not every population size was simulated to save on time


AT_standard_traitsd <- equilibriumpopulationsizeGamma(mu = mu, sigma = testvec, w = w) #Calculating equilibrium population size for the used values for sigma

par(mar = c(5.1, 2.5, 4.1, 0.1))

#Plot for the different values for the trait sd

matplot(startingpop, mate_finding_trait_sd[[1]],type = "l", xlab = "Founder population size", ylab = "",col = "#011C40FF", cex.axis = 1.8, cex.lab = 1.8, lwd = 3, ylim = c(0,1), main = "B", cex.main = 1.8)
matlines(startingpop, mate_finding_trait_sd[[2]],col = "#40e078", lwd = 3)
matlines(startingpop, mate_finding_trait_sd[[3]], col = "#05DBF2FF", lwd = 3)
matlines(startingpop, mate_finding_trait_sd[[4]], col = "#4C4CFFFF", lwd = 3)
abline(v = AT_standard_traitsd, col = "#40e078", lwd = 3, lty = c(1,2,3))
abline(v = AT_none, col = "#4C4CFFFF", lwd = 3)
legend("topleft", title = "Trait sd:",  legend = c(paste(testvec)), col = c("black", "black", "black"), lty = c(1,2,3), lwd = rep(2,3), cex = 1.6 , bty="o")

par(mar = c(5.1, 2.5, 4.1, 1))

#Plot for the different values for the mutation sd

matplot(startingpop, mate_finding_mutation_sd[[1]],type = "l", xlab = "Founder population size", ylab = "",col = "#011C40FF", cex.axis = 1.8, cex.lab = 1.8, lwd = 3, ylim = c(0,1), main = "C", cex.main = 1.8)
matlines(startingpop, mate_finding_mutation_sd[[2]],col = "#40e078", lwd = 3)
matlines(startingpop, mate_finding_mutation_sd[[3]], col = "#05DBF2FF", lwd = 3)
matlines(startingpop, mate_finding_mutation_sd[[4]], col = "#4C4CFFFF", lwd = 3)
abline(v = AT_standard, col = "#40e078", lwd = 3)
abline(v = AT_none, col = "#4C4CFFFF", lwd = 3)
legend("topleft", title = "Mutation sd:",  legend = c(paste(testvec)), col = c("black", "black", "black"), lty = c(1,2,3), lwd = rep(2,3), cex = 1.6 , bty="o")

par(mfrow=c(1,1))

dev.print(dev=pdf,"mate_finding_mainresults.pdf")

####Making the plot for predator driven Allee effect

load("Simulation_results/Predator_driven/Pred_prey_standard_all_population_sizes.RData")

load("Simulation_results/Predator_driven/Pred_prey_different_mutation_sd_500_replicates_0.001_0.005_0.01_pop_sequence_10_800_10_Schritte.RData")

load("Simulation_results/Predator_driven/Pred_prey_different_trait_sd_500_replicates_0.001_0.005_0.01_pop_sequence_10_800_10_Schritte.RData")

pred_prey_standard <- Pred_prey_standard_all_population_sizes

pred_prey_mutation_sd <- Pred_prey_different_mutation_sd_500_replicates_0.001_0.005_0.01_pop_sequence_10_800_10_Schritte

pred_prey_trait_sd <- Pred_prey_different_trait_sd_500_replicates_0.001_0.005_0.01_pop_sequence_10_800_10_Schritte


#Equation for the equilibrium population size (also covers scenario where sigma is 0)

equilibriumpopulationsizeGamma_PP <- function(mu, sigma, w, b, c, h,P){
  A <- (mu/(b*sigma^2+mu))^(mu^2/sigma^2)
  if (sigma == 0){
    Nstar <- w*P/(h*(w-2)) - 1/(h*c*exp(-mu*b))
  } else {
    Nstar <- w*P/(h*(w-2)) - 1/(h*c*A)
  }
  return(Nstar)
}

#Setting standard parameters used in the model for calculating the equilibrium population size


mu = 0.02
sigma = 0.005
w = 2.1 
b = 70
c = 2
h = 0.03
P = 1

#Calculating equilibrium population size for the standard value of sigma and if sigma is 0


AT_standard_PP <- equilibriumpopulationsizeGamma_PP(mu = mu, sigma = sigma, w = w,  b = b, c = c, h = h, P = P)
AT_none_PP <- equilibriumpopulationsizeGamma_PP(mu = mu, sigma = 0, w = w,  b = b, c = c, h = h, P = P)


#Making the plots for the predator driven Allee effect

par(mfrow=c(1,3))
par(mar = c(5.1, 4.5, 4.1, 0.1))

startingpop <- seq(10,800,1) #Starting population sizes

testvec <-c(0.001,0.005,0.01) #Values used when testing different values for the trait and mutation sd 

#Plot for the standard values

matplot(startingpop, pred_prey_standard[[1]],type = "l", xlab = "Founder population size", ylab = "Proportion of surviving populations",col = "#011C40FF", cex.axis = 1.8, cex.lab = 1.8, lwd = 3, ylim = c(0,1), main = "A", cex.main = 1.8)
matlines(startingpop, pred_prey_standard[[2]],col = "#40e078", lwd = 3)
matlines(startingpop, pred_prey_standard[[3]], col = "#05DBF2FF", lwd = 3)
matlines(startingpop, pred_prey_standard[[4]], col = "#4C4CFFFF", lwd = 3)
abline(v = AT_standard_PP, col = "#40e078", lwd = 3)
abline(v = AT_none_PP, col = "#4C4CFFFF", lwd = 3)
legend("topleft", title = "Model includes:", legend = c("ITV + Inheritance", "ITV only", "Inheritance only", "None"), col = c("#011C40FF", "#40e078", "#05DBF2FF", "#4C4CFFFF"), lty = rep(1,5), lwd = c(rep(2,4)) ,ncol = 1, cex = 1.6 , bty="o")


startingpop <- seq(10,800,10) #For these graphs not every population size was simulated to safe on time

#Calculating equilibrium population size for the used values for sigma

AT_standard_traitsd_PP <- numeric(length(testvec))

for (i in 1:length(AT_standard_traitsd_PP)) {
  
  AT_standard_traitsd_PP[i] <- equilibriumpopulationsizeGamma_PP(mu = mu, sigma = testvec[i], w = w,  b = b, c = c, h = h, P = P)
  
  
}

par(mar = c(5.1, 2.5, 4.1, 0.1))

#Plot for the different values for the trait sd

matplot(startingpop, pred_prey_trait_sd[[1]],type = "l", xlab = "Founder population size", ylab = "",col = "#011C40FF", cex.axis = 1.8, cex.lab = 1.8, lwd = 3, ylim = c(0,1), main = "B", cex.main = 1.8)
matlines(startingpop, pred_prey_trait_sd[[2]],col = "#40e078", lwd = 3)
matlines(startingpop, pred_prey_trait_sd[[3]], col = "#05DBF2FF", lwd = 3)
matlines(startingpop, pred_prey_trait_sd[[4]], col = "#4C4CFFFF", lwd = 3)
abline(v = AT_standard_traitsd_PP, col = "#40e078", lwd = 3, lty = c(1,2,3))
abline(v = AT_none_PP, col = "#4C4CFFFF", lwd = 3)
legend("topleft", title = "Trait sd:",  legend = c(paste(testvec)), col = c("black", "black", "black"), lty = c(1,2,3), lwd = rep(2,3), cex = 1.6 , bty="o")

par(mar = c(5.1, 2.5, 4.1, 1))

#Plot for the different values for the mutation sd

matplot(startingpop, pred_prey_mutation_sd[[1]],type = "l", xlab = "Founder population size", ylab = "",col = "#011C40FF", cex.axis = 1.8, cex.lab = 1.8, lwd = 3, ylim = c(0,1), main = "C", cex.main = 1.8)
matlines(startingpop, pred_prey_mutation_sd[[2]],col = "#40e078", lwd = 3)
matlines(startingpop, pred_prey_mutation_sd[[3]], col = "#05DBF2FF", lwd = 3)
matlines(startingpop, pred_prey_mutation_sd[[4]], col = "#4C4CFFFF", lwd = 3)
abline(v = AT_standard_PP, col = "#40e078", lwd = 3)
abline(v = AT_none_PP, col = "#4C4CFFFF", lwd = 3)
legend("topleft", title = "Mutation sd:",  legend = c(paste(testvec)), col = c("black", "black", "black"), lty = c(1,2,3), lwd = rep(2,3), cex = 1.6 , bty="o")


par(mfrow=c(1,1))

dev.print(dev=pdf,"predator_prey_mainresults.pdf")
```
