## Supplementary material for "How does individual trait variation impact the survival of populations with an Allee effect?": R code and simulation data: Fig4_FigS2.R

```
source("All_functions_predator_prey.R")
results0.001 <- alleefunc_pred(mutation_sd=0.001,replicates=500)
results0.005 <- alleefunc_pred(mutation_sd=0.005,replicates=500)
results0.01 <- alleefunc_pred(mutation_sd=0.01,replicates=500)

sum(results0.001[[3]])/500
sum(results0.005[[3]])/500
sum(results0.01[[3]])/500

sd0.001 <- rowMeans(results0.001[[4]],na.rm=T)
sd0.005 <- rowMeans(results0.005[[4]],na.rm=T)
sd0.01 <- rowMeans(results0.01[[4]],na.rm=T)

means0.001 <- rowMeans(results0.001[[2]],na.rm=T)
means0.005 <- rowMeans(results0.005[[2]],na.rm=T)
means0.01 <- rowMeans(results0.01[[2]],na.rm=T)

popsize0.001 <- rowMeans(results0.001[[1]],na.rm=T)
popsize0.005 <- rowMeans(results0.005[[1]],na.rm=T)
popsize0.01 <- rowMeans(results0.01[[1]],na.rm=T)

#Make Fig. 4
par(mfrow=c(1,3))
matplot(0:100,cbind(popsize0.001,popsize0.005,popsize0.01),xlab="Generation",ylab="Population size",type="l",col="black",lty=c(1:3),main="A")
legend("topleft",title="Mutation sd",legend=c(0.001,0.005,0.01),lty=1:3)
matplot(0:100,cbind(means0.001,means0.005,means0.01),xlab="Generation",ylab="Trait mean",type="l",col="black",lty=c(1:3),main="B")
matplot(0:100,cbind(sd0.001,sd0.005,sd0.01),xlab="Generation",ylab="Trait standard deviation",type="l",col="black",lty=c(1:3),main="C")
dev.print(dev=pdf,"timeseries_predatordriven.pdf")

#Make Fig. S2
par(mfrow=c(1,1))
popsizes0.01 <- results0.01[[1]]
matplot(popsizes0.01[,1:30],xlab="Generations",ylab="Population size",type="l",col=rainbow(30))
dev.print(dev=pdf,"populationsizes_predatordriven_sdmut0_01.pdf")
```
