## Supplementary material for "How does individual trait variation impact the survival of populations with an Allee effect?": R code and simulation data: Fig5.R

```
###Plot for the estimate of the effect of ITV for the mate finding Allee effect (Fig. 5)

par(mfrow=c(1,2))

#A random trait value is drawn for a set number of (female) individuals and their mating probability is calculated. Their average mating probability is compared to the mating probability resulting from the mean trait value 

mating <- function(M,r){ #Function to calculate the mating probability for an individual with the given trait value r
  1-exp(-r*M)
}

mean_r <- 0.02 #Mean trait value
trait_sd <- 0.005 #trait standard deviation value
num_test <- 1000 #Number of tested (female) individuals
r <- rgamma(n = num_test, shape = (mean_r/trait_sd)^2 ,rate = mean_r/trait_sd^2) #drawing a trait value of each tested (female) individual
m <- 100 #number of males 

matingvec <- mating(m,r) #Calculating the mating probability of each female based on their trait value and the set number of males  
value_mate_finding <- round((mean(matingvec)-mating(m,mean(r)))/mating(m,mean(r)),digits = 3) #Calculating the estimate of the effect of ITV

#Plotting the results

plot(r,matingvec, xlab = "Trait values", ylab = "Resulting mating probability", ylim = c(0.5, 1), cex.axis = 1.4, cex.lab = 1.4, main = "A")
points(mean(r),mean(matingvec),col ="red",pch = 19,cex = 2)
points(mean(r),mating(m,mean(r)),col ="purple",pch = 19,cex =2 )
legend("topleft",legend = c(expression(bar(f(x))),expression(f(bar(x)))),pch = 19,col = c("red","purple"),cex = 1.4)
text(cex = 1.4, mean_r+0.004,0.55,bquote(paste("Estimate of the effect of ITV"," = ",frac(bar(f(x))-f(bar(x)),f(bar(x)))," = ",.(value_mate_finding))))


###Plot for Predator prey

#A random trait value is drawn for a set number of individuals and their survival probability is calculated. Their average survival probability is compared to the survival probability resulting from the mean trait value 

survival_prob <- function(P,c,b,t,h,trait_vec){ #Function to calculate the survival probability of each tested individual
  1-((c*exp(-b*t))/(1+sum(c*exp(-b*trait_vec)*h))*P)
}

#Set parameters used in the equation

N <- 100 #Number of individuals that make up the rest of the population  
P <- 1 #Number of predators
c <- 2 #Maximum attack rate of the predator
b <- 70 #Decay rate of the attack rate with increasing trait
mean_t <- 0.02 #Mean trait value
trait_sd <- 0.005 #Trait standard deviation value
num_test <- 1000 #Number of tested individuals  
t <- rgamma(n = num_test, shape = (mean_t/trait_sd)^2 ,rate = mean_t/trait_sd^2) #Generate trait value for each tested individual
h <- 0.03 #Handling time of the predator

survival_probvec <- numeric(length(t))

#Calculate the survival probability for each tested individual

for (k in 1:length(t)) {
  trait_vec <- rgamma(n = N, shape = (mean_t/trait_sd)^2 ,rate = mean_t/trait_sd^2) #Generate trait value for each individual in the rest of the simulated population
  
  survival_probvec[k] <- survival_prob(P,c,b,t[k],h,trait_vec) #Calculate the survival probability for each tested individual based on the trait values of the individuals that make up the rest of the population
}

#Calculate the estimate of the effect of ITV
value_pred_prey <- round((mean(survival_probvec)-survival_prob(P,c,b,mean(t),h,rep(mean_t,N)))/survival_prob(P,c,b,mean(t),h,rep(mean_t,N)),digits = 3)

#Plotting the results

plot(t,survival_probvec, xlab = "Trait values", ylab = "Resulting survival probability", ylim = c(0.5, 1), cex.axis = 1.4, cex.lab = 1.4, main = "B")
points(mean(t),mean(survival_probvec),col ="red",pch = 19,cex = 2)
points(mean(t),survival_prob(P,c,b,mean(t),h,rep(mean_t,N)),col ="purple",pch = 19,cex =2 )
legend("topleft",legend = c(expression(bar(f(x))),expression(f(bar(x)))),pch = 19,col = c("red","purple"),cex = 1.4)
text(cex = 1.4, c(mean_t+0.004),0.55,bquote(paste("Estimate of the effect of ITV"," = ",frac(bar(f(x))-f(bar(x)),f(bar(x)))," = ",.(value_pred_prey))))


par(mfrow=c(1,1))
```
