## Supplementary material for "How does individual trait variation impact the survival of populations with an Allee effect?": R code and simulation data: generate_results_mate_finding.R

```
#Generating the results for the mate finding Allee effect model

library(foreach)
library(doParallel)

source("All_functions_mate_finding.R")

cores <- detectCores() #Detecting available cores

cl <- makeCluster(cores-4) #Setting how many cores to use

registerDoParallel(cl) 


startingpop <- seq(10,800,10) #Vector for the starting populations
testvec <-c(0.001,0.005,0.01) #Vector used to set the values for either the trait standard variation or mutation standard variation  
replicates <- 500 #Number of replicates per run

results <- matrix(nrow = length(startingpop),ncol = length(testvec)) #Making empty matrix to save the results in

Results_list <- foreach (i = 1:4) %dopar% { #Running the four different model versions at the same time
  
  if(i == 1){
    
    for (r in 1:length(testvec)){ #looping over the three different values for the trait sd or mutation sd
      
      for (p in 1:length(startingpop)){ #looping over every starting population
        
        results[p,r]<-mean(alleefunc(N0 = startingpop[p], mutation_sd = testvec[r], replicates = replicates)[[3]])
        
      }
    }
  }
  
  if(i == 2){
    
    for (r in 1:length(testvec)) {
      
      for (p in 1:length(startingpop)){
        
        results[p,r]<-mean(noinheritance(N0 = startingpop[p], mutation_sd = testvec[r], replicates = replicates)[[3]])
        
      }
    }
  }
  
  if(i == 3){
    
    for (r in 1:length(testvec)) {
      
      for (p in 1:length(startingpop)) {
        
        results[p,r]<-mean(noITV(N0 = startingpop[p], mutation_sd = testvec[r], replicates = replicates)[[3]])
        
      }
    }
  }
  
  if(i == 4){
    
    for (r in 1:length(testvec)) {
      
      for (p in 1:length(startingpop)) {
        
        results[p,r]<-mean(noinheritance_ITV(N0 = startingpop[p], mutation_sd = testvec[r], replicates = replicates)[[3]])
        
      }
    }
  }
  results
}

#Code was run three different times. Once with changing the mutation standard deviation values, once with changing the trait standard devitaion and once with keeping both values as the standard
#To run the model for different trait sd values the mutation_sd was changed to traitsd
#To run the model with just the standard values for the trait sd and mutation sd the testvect and the loop over the testvec were removed.
#All results were saved as .RData files (see folder Simulation_results).

stopCluster(cl) #Stop the cluster
```
