## Supplementary material for "How does individual trait variation impact the survival of populations with an Allee effect?": R code and simulation data: README.txt

This folder contains the R code and main simulation results (those that take longer to run) for the manuscript "How does individual trait variation impact the survival of populations with an Allee effect?".

The files All_functions_mate_finding.R and All_functions_predator_prey.R contain the main simulation functions for the different model versions of the mate-finding Allee effect model and the predator-driven Allee effect model.

The functions generate_results_mate_finding.R and generate_results_predator_prey.R generate the main results underlying Fig. 1 and Fig. 3, respectively. Note that running them takes some time, but we are providing the simulation results in the folder Simulation_results.

The other R scripts are for generating the various figures.
