## Supplementary material for "How does individual trait variation impact the survival of populations with an Allee effect?": Maxima code for analytical apprimation: supplement_maxima.html


Please enable JavaScript in order to get a 2d display of the equations embedded in this web page.

\( \DeclareMathOperator{\abs}{abs}
\newcommand{\ensuremath}[1]{\mbox{$#1$}}
\)

Supplementary Material 1: Maxima file supporting the analytical calculation of the Allee threshold

1 Setting up the Gamma distribution

Choose the parameters of the Gamma distribution to obtain a desired mean and standard deviation:

|  |  |
| --- | --- |
| (%i1) | alpha: mu^2/sigma^2; |

\[\]\[\tag{lph} \frac{{{mu}^{2}}}{{{sigma}^{2}}}\]

|  |  |
| --- | --- |
| (%i2) | lambda: mu/sigma^2; |

\[\]\[\tag{ambd} \frac{mu}{{{sigma}^{2}}}\]

|  |  |
| --- | --- |
| (%i3) | assume(sigma>0); |

\[\]\[\tag{%o3} \left[ sigma\mathop{> }0\right] \]

|  |  |
| --- | --- |
| (%i4) | assume(mu>0); |

\[\]\[\tag{%o4} \left[ mu\mathop{> }0\right] \]

|  |  |
| --- | --- |
| (%i5) | assume(w>2); |

\[\]\[\tag{%o5} \left[ w\mathop{> }2\right] \]

|  |  |
| --- | --- |
| (%i6) | gammapdf(z):= (lambda^alpha)/gamma(alpha) · z^(alpha−1) ·exp(−lambda·z) ; |

\[\]\[\tag{%o6} \mathop{gammapdf}(z)\mathop{:=}\frac{{{lambda}^{alpha}}}{\mathop{gamma}(alpha)} {{z}^{alpha\mathop{-}1}} \mathop{exp}\left( \mathop{-}lambda z\right) \]

2 Mate-finding Allee effect

|  |  |
| --- | --- |
| (%i7) | assume(N>0); |

\[\]\[\tag{%o7} \left[ N\mathop{> }0\right] \]

|  |  |
| --- | --- |
| (%i8) | averagematingsuccess: ratsimp(integrate((1−exp(−r·N/2))·gammapdf(r),r,0,inf)); |

\[\]\[\mbox{}\\"Is "\frac{{{mu}^{2}}}{{{sigma}^{2}}}" an "\ensuremath{\mathrm{integer}}"?"no;\]

\[\]\[\tag{veragematingsucces} \mathop{-}\left( \frac{{{mu}^{\frac{{{mu}^{2}}}{{{sigma}^{2}}}}} {{2}^{\frac{{{mu}^{2}}}{{{sigma}^{2}}}}}\mathop{-}{{\left( N {{sigma}^{2}}\mathop{+}2 mu\right) }^{\frac{{{mu}^{2}}}{{{sigma}^{2}}}}}}{{{\left( N {{sigma}^{2}}\mathop{+}2 mu\right) }^{\frac{{{mu}^{2}}}{{{sigma}^{2}}}}}}\right) \]

Solve for the Allee threshold:

|  |  |
| --- | --- |
| (%i9) | solve([w·averagematingsuccess = 2],[N]); |

\[\]\[\mbox{}\\"Is "\frac{{{mu}^{2}}}{{{sigma}^{2}}}" an "\ensuremath{\mathrm{integer}}"?"no;\]

\[\]\[\tag{%o9} \left[ N\mathop{=}\frac{2 mu {{w}^{\frac{{{sigma}^{2}}}{{{mu}^{2}}}}}\mathop{-}2 mu {{\left( w\mathop{-}2\right) }^{\frac{{{sigma}^{2}}}{{{mu}^{2}}}}}}{{{sigma}^{2}} {{\left( w\mathop{-}2\right) }^{\frac{{{sigma}^{2}}}{{{mu}^{2}}}}}}\right] \]

3 Predator-driven Allee effect

|  |  |
| --- | --- |
| (%i10) | assume(c>0); |

\[\]\[\tag{%o10} \left[ c\mathop{> }0\right] \]

|  |  |
| --- | --- |
| (%i11) | assume(h>0); |

\[\]\[\tag{%o11} \left[ h\mathop{> }0\right] \]

|  |  |
| --- | --- |
| (%i12) | assume(P>0); |

\[\]\[\tag{%o12} \left[ P\mathop{> }0\right] \]

|  |  |
| --- | --- |
| (%i13) | assume(b>0); |

\[\]\[\tag{%o13} \left[ b\mathop{> }0\right] \]

|  |  |
| --- | --- |
| (%i14) | A: ratsimp(integrate(exp(−b·z)·gammapdf(z),z,0,inf)); |

\[\]\[\mbox{}\\"Is "\frac{{{mu}^{2}}}{{{sigma}^{2}}}" an "\ensuremath{\mathrm{integer}}"?"no;\]

\[\]\[\tag{} \frac{{{mu}^{\frac{{{mu}^{2}}}{{{sigma}^{2}}}}}}{{{\left( b {{sigma}^{2}}\mathop{+}mu\right) }^{\frac{{{mu}^{2}}}{{{sigma}^{2}}}}}}\]

---

Created with wxMaxima.

 The source of this Maxima session can be downloaded here.
